# Exploring integument transcriptomes, cuticle ultrastructure, and cuticular hydrocarbons profiles in eusocial and solitary bee species displaying heterochronic adult cuticle maturation

**DOI:** 10.1101/437962

**Authors:** Tiago Falcon, Daniel G. Pinheiro, Maria Juliana Ferreira-Caliman, Izabel C. C. Turatti, Fabiano C. Pinto de Abreu, Juliana S. Galaschi-Teixeira, Juliana R. Martins, Moysés Elias-Neto, Michelle P. M. Soares, Michelle P. M. Soares, Marcela B. Laure, Vera L. C. Figueiredo, Norberto Peporine Lopes, Zilá L. P. Simões, Carlos A. Garófalo, Márcia M. G. Bitondi

## Abstract

Differences in the timing of exoskeleton melanization and sclerotization are evident when comparing eusocial and solitary bees. This cuticular maturation heterochrony may be associated with life style, considering that eusocial bees remain protected inside the nest for many days after emergence, while the solitary bees immediately start outside activities. To address this issue, we characterized gene expression using large-scale RNA sequencing (RNA-seq), and quantified cuticular hydrocarbon (CHC) through gas chromatography-mass spectrometry in comparative studies of the integument (cuticle plus its underlying epidermis) of two eusocial and a solitary bee species. In addition, we used transmission electron microscopy (TEM) for studying the developing cuticle of these and other three bee species also differing in life style. We found 13,200, 55,209 and 30,161 transcript types in the integument of the eusocial *Apis mellifera* and *Frieseomelitta varia*, and the solitary *Centris analis*, respectively. In general, structural cuticle proteins and chitin-related genes were upregulated in pharate-adults and newly-emerged bees whereas transcripts for odorant binding proteins, cytochrome P450 and antioxidant proteins were overrepresented in foragers. Consistent with our hypothesis, a distance correlation analysis based on the differentially expressed genes suggested delayed cuticle maturation in *A. mellifera* in comparison to the solitary bee. However, this was not confirmed in the comparison with *F. varia*. The expression profiles of 27 of 119 genes displaying functional attributes related to cuticle formation/differentiation were positively correlated between *A. mellifera* and *F. varia*, and negatively or non-correlated with *C. analis*, suggesting roles in cuticular maturation heterochrony. However, we also found transcript profiles positively correlated between each one of the eusocial species and *C. analis*. Gene co-expression networks greatly differed between the bee species, but we identified common gene interactions exclusively between the eusocial species. Except for *F. varia*, the TEM analysis is consistent with cuticle development timing adapted to the social or solitary life style. In support to our hypothesis, the absolute quantities of n-alkanes and unsaturated CHCs were significantly higher in foragers than in the earlier developmental phases of the eusocial bees, but did not discriminate newly-emerged from foragers in *C. analis*. By highlighting differences in integument gene expression, cuticle ultrastructure, and CHC profiles between eusocial and solitary bees, our data provided insights into the process of heterochronic cuticle maturation associated to the way of life.

**Author Summary:** From our previous observation that bees with distinct habits of life, eusocial and solitary, exhibit different degrees of cuticle melanization and sclerotization at the emergence, we decided to analyze the genetic signatures and ultrastructure of the integument, as well as the CHC profiles that could be involved in cuticle maturation. The expression patterns of certain genes involved in the melanization/sclerotization pathway, chitin metabolism, cuticle structure, and also regulators of cuticle renewal and tanning, in addition to other genes, might be grounded the slow process of cuticle maturation in the eusocial bees in comparison to the solitary ones. The electron micrographs revealed differences in the timing of cuticle deposition for the eusocial and solitary species. Among the identified CHCs, the proportions and quantities of n-alkanes in the developing cuticle are consistent with the faster cuticular maturation in the solitary bee, thus supporting our hypothesis.

## Introduction

The exoskeleton (cuticle) enables arthropods to exploit a multitude of ecological habitats, and is central to the evolutionary success and worldwide expansion of insects. It is necessary for muscles attachment, for protection against predators, injuries, and pathogens [1]. In addition, its thickness is positively correlated with the resistance to some types of insecticides [2]. The exoskeleton is periodically shed and a new, larger one is formed, this characterizing the successive molting episodes that allow for insect growth and development. Its composition is defined by the secretion of products synthesized by the epidermis as well as by the uptake of molecules from other sources, for instances, hemolymph [3]. These products are used for cuticle renewal at each molting episode coordinated by changes in the titer of 20-hydroxyecdysone (20E), the active product of ecdysone hydroxylation. The Ashburner model postulated to explain 20E-induced chromosomal puffs in the larval salivary glands of *D. melanogaster* have ultimately led to the knowledge of molecular elements regulating molting and metamorphosis [4]. When 20E binds to the heterodimeric receptor consisting of EcR (Ecdysone receptor) and Usp (Ultraspiracle) proteins, its trigger a transcription factor regulatory cascade. Upstream elements of this cascade respond to the high 20E titer that also induces apolysis and initiates molting, whereas most downstream elements are only induced by the subsequent decrease in 20E titer. Binding sites for several of the transcription factors in this cascade were identified in many cuticular protein genes [5], suggesting that they, and other genes involved in cuticle remodeling [6, 7] are indirectly regulated by 20E.

The exoskeleton comprises an inner procuticle formed by layers of endocuticle and exocuticle, an outer epicuticle and the superficial envelope. The procuticle consists of a variety of proteins and chitin, a polymer of the glucose-derived N-acetylglucosamine. Chitin is a major compound in the insect exoskeleton [8]. Key enzymes in the chitin biosynthetic pathway starting from trehalose are the highly conserved chitin synthases that catalyze the transformation of UDP-*N*-acetylglucosamine to chitin. Chitin-modifying enzymes, specifically chitin deacetylases (Cdas), catalyze the conversion of chitin to chitosan, a polymer of *β*-1,4-linked d-glucosamine residues. Mutations in Cda genes are lethal to insect embryos, suggesting that these enzymes play critical roles during development, including the molting process [9]. Molting involves digestion of the actual cuticle, a process mediated by chitin-degrading en-zymes, chitinases, which accumulate in the molting fluid [10]. The epicuticle does not contain chitin, but contains proteins and lipids and is rich in quinones, which are oxidized derivatives of aromatic compounds [11]. Together with chitin, the structural cuticular proteins constitute the bulk of insect cuti-cle. Based on defining sequence domains, they have been classified into twelve families [12]. Proteins in the CPR family, with the largest number of members, contain the R&R Consensus [13, 14]. Some other structural cuticular proteins pertain to the Tweedle (Twdl) class [15], or were classified as Cuticular Proteins of Low Complexity – Proline-rich (CPLCP), Cuticular Proteins with Forty-four amino acid residues (CPF), Cuticular proteins analogous to peritrophins (Cpap), Glycine-Rich cuticular Proteins (GRP), and apidermins, among other classes. Some cuticular proteins, however, do not fill the features for inclusion in the pre-established classes. The main components of the envelope are the cuticular hydrocarbons (CHC) [16] that play roles in chemical communication (unsaturated CHC) [17, 18] and, together with other lipids, act as a barrier against insect desiccation by preventing water loss (mainly n-alkanes) [19, 17]. Key enzymes in CHC biosynthetic pathways occurring in the epidermis-associated oenocytes are the desaturases and elongases [20–22]. We previously determined gene expression profiles of six desaturases and ten elongases in the developing integument of *A. mellifera*, and correlated them with n-alkanes, methyl-alkanes, dimethyl-alkanes, alkenes and alkadienes quantification profiles [23]. Besides highlighting the CHC composition underlying envelope formation, these data provided clues to predict the function of these genes in CHC biosynthetic pathways.

In addition to chitin, cuticular proteins, CHCs, and other compounds, melanin pigments are crucial for the exoskeleton formation in insects. The chemical reactions in the core of the melanin biosynthetic pathway are evolutionary conserved. This pathway comprises the conversion of tyrosine into 3,4-dihydroxyphenylalanine (dopa) by the action of tyrosine hydroxylase (TH). Dopa is converted to dopamine, the primary precursor of insect melanin, via a decarboxylation reaction catalyzed by dopa decarboxylase (Ddc). Dopa or dopamine is further oxidized to dopaquinone or dopaminequinone, and finally these pigment precursors are converted into dopa-melanin or dopamine-melanin through reactions catalyzed by dopachrome conversion enzyme, a product of the *yellow* gene, and laccase2. Alternatively, dopamine is acetylated to N-acetyl-dopamine (NADA), and in conjugation with α-alanine originates N-β-alanyldopamine (NBAD). Both catechols are precursors for production of colorless and yellowish sclerotins [24, 25]. Thus, melanization occurs concomitantly to sclerotization through a shared biosynthetic pathway. Both processes are fundamental for the exoskeleton development [26], and are developmentally regulated by 20E [27, 28].

Among bees, we can distinguish the solitary and eusocial species. In the solitary species, every female constructs its own nest where it lay eggs, but does not provide care for the ecloded larvae. In contrast, the social organization is grounded on the division of labor between fertile queens and more or less sterile, or completely sterile, workers that are engaged in nest construction and maintenance, besides caring for the queen’s offspring [29, 30]. The search for genomic signatures of eusociality evolution in bees has grown since the publication of the *A. mellifera* genome [31] and gained force with the recent release of two *Bombus* species genomes [32] and the study of Kapheim *et al.* [33] comparing the genomes of ten bee species.

In this context, we draw our attention to the fact that bees greatly vary in the grade of cuticle melanization/sclerotization at the emergence time (adult ecdysis). In a previous study on the morphology of the developing adult cuticle [34], we observed that in eusocial bees, but not in the solitary ones, the process of cuticle melanization/sclerotization leading to cuticle maturation is extended to the adult stage. After emergence, workers from eusocial species (including the primitively eusocial bees from Bombini) spend some days performing inside nest activities, and during this period they stay protected in a safe and provisioned environment [35] where the hygienic behavior provides a certain level of immunity [36]. In contrast, the newly emerged solitary bees immediately leave the nest. Therefore, they need a fully mature cuticle to protect them in the external environment. This shift in the timing of cuticle maturation seems a case of heterochrony, which is defined as a change in the timing of development of a tissue or anatomical part relative to an ancestor, or between taxa [37]. If this assumption proves to be true, it can entail a link between the rate of cuticle maturation and the evolution of sociality in insects.

Here, we used the integument (cuticle and its subjacent epidermis) in an approach based on large-scale RNA sequencing (RNA-seq), transmission electron microscopy (TEM) and gas chromatography-mass spectrometry (GC/MS) to describe cuticle maturation in two eusocial bee species, *Apis mellifera* (Apini) and *Frieseomelitta varia* (Meliponini), and a solitary bee species, *Centris analis* (Centridini), thesolitarylifestylebeingconsideredtheancestralconditionforbes[38]. TEM was also used for studying the ultrastructure of the cuticle of the primitively eusocial bee, *Bombus brasiliensis* (Bombini), the facultatively eusocial *Euglossa cordata* (Euglossini), and the solitary bee, *Tetrapedia diversipes*. This combined approach allowed us to compare the integuments at the morphological and molecular levels, besides highlighting differences that could be related to the heterochronic process of cuticle maturation. Among the genes expressed in the integument, we focused on those involved in the melanization/sclerotization pathway, chitin metabolism, genes encoding structural cuticular proteins, regulators of cuticle renewal and tanning, desaturase and elongase genes potentially involved in CHC biosynthesis, circadian clock genes that could determine the rhythm of cuticle layers deposition [39, 40], and genes encoding pigments other than melanin.

The comparison of integument transcriptomes of three bee species at developmental points corresponding to adult cuticle formation, ecdysis, and at a mature age (foragers) gave us back the discovery of distinct genetic signatures of the integument, and highlighted differences in gene set expression profiles. The use of TEM and CHC analysis complemented these data by adding new information on cuticle ultrastructure and chemical profiles of its superficial layer, the envelope.

## Results

### Differential gene expression in the integument of *A. mellifera*, *F. varia* and *C. analis* during adult cuticle formation/maturation

We identified the expression of 13,200 genes in the developing integument of *A. mellifera*, and 55,209 and 30,161 contigs in the developing integument of *F. varia* and *C. analis*, respectively (S1 File). The data obtained from the three biological samples of each developmental phase, Pbm (pharate adult), Ne (newly-emerged) and Fg (forager) of each bee species, in a total of 27 transcriptomes, were used in Pearson correlation analysis in order to check reproducibility. A hierarchical clustering on pairwise correlation is shown in S1 Fig. In general, the samples of the same developmental phase (biological triplicates) joined together, indicating that they are more similar to each other than to samples of the other developmental phases. As expected, for the three bee species, the least correlated samples were those originated from the Pbm and Fg integuments. When filtering these data sets for the genes (DEGs) or contigs (DECs) differentially expressed between the developmental phases, we found 3,184 DEGs for *A. mellifera*, 5,959 DECs for *F. varia* and 2,543 DECs for *C. analis*, representing 24.1%, 10.8%, and 8.4% of the identified genes, respectively. Fig 1 shows the number of genes that were upregulated in the comparisons between the developmental phases of each of the three bee species. In *A. mellifera*, 14.8% and 17.8% of the DEGs were upregulated in the Pbm phase in comparison to the Ne and Fg phases, respectively; 20.9% and 7.8% were upregulated in Ne in comparison to Pbm and Fg; 24.6% and 10.4% were more expressed in Fg than Pbm and Ne. In *F. varia*, 21.1% and 39.3% of the DECs were upregulated in Pbm compared to the Ne and Fg phases, respectively; 27.9% and 21.1% DECs were more expressed in Ne than in Pbm and Fg; 39.6% and 16.0% showed higher expression in Fg than in Pbm and Ne. In *C. analis*, the Pbm phase showed higher expression for 39.2% and 31.1% of the DECs in comparison to the Ne and Fg phases; 32.7% and 6.1% of the DECs were upregulated in Ne in comparison to Pbm and Fg; 31.9% and 3.6% were more expressed in Fg than in Pbm and Ne. These proportions of upregulated genes significantly vary in the comparisons of the developmental phases of each bee species and also between the bee species. In addition, the proportions of genes upregulated in the adult phases (Ne *versus* Fg) were significantly lower in the solitary *C. analis* than in the eusocial *A. mellifera* and *F. varia* bee species (z test, p ≤ 0.001, except for one of the comparisons where p = 0.014).

**Fig 1.**
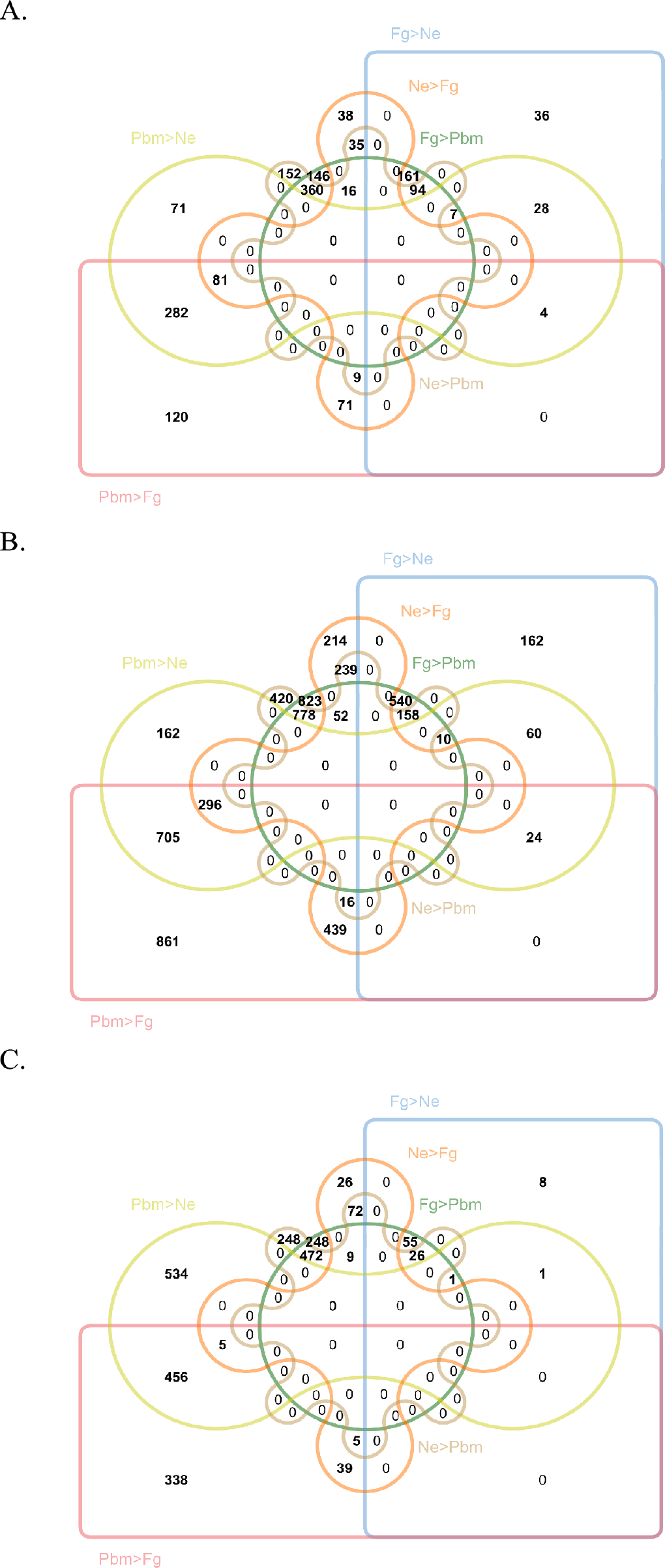
Venn diagrams constructed with the genes and contigs differentially expressed in the integument of the developmental phases of (A) *A. mellifera*, (B)*F. varia*, and (C) *C. analis*. The number of genes upregulated in each pairwise comparison is indicated. Pbm: pharate adults; Ne: newly-emerged bees; Fg: foragers.

To make more comprehensive the RNA-seq analysis of the integument, we searched the Gene Ontology (GO) functional terms for all *A. mellifera* DEGs and all *F. varia* and *C. analis* DECs. The GO annotations for Molecular Function, Cellular Component and Biological Process categories are described in S3 File. We then extracted from this analysis the functional terms more evidently related to cuticle development (Fig 2). Structural molecule activity, chitin-binding, and chitin metabolic process were categories overrepresented in the younger phases, i.e., the Pbm and Ne phases of the three bee species. Structural constituent of cuticle, structural constituent of chitin-based cuticle, and other cuticular components-related GO categories also included genes more expressed in the Pbm and Ne integuments of both, or one of the eusocial species. Functional categories related to the epidermis, which is the tissue responsible for secreting the cuticle, specifically epithelium development, epithelial cell differentiation/development, cell adhesion, cell junction organization/assembly, among other categories, were also more represented in the younger Pbm and Ne bees, but only of the eusocial species. For the three bee species, the DEGs and DECs more related to the functionality of the integument of newly-emerged (Ne) and forager bees (Fg) (here named older phases for simplification), were included in the following overrepresented GO terms: fatty acid biosynthetic process, lipid biosynthetic process, organic acid biosynthetic process, and carboxylic acid biosynthetic process. These terms and others overrepresented in the older Ne and Fg phases of *F. varia* and *C. analis*, i.e., very-long-chain fatty acid metabolic process, and fatty acid metabolic process, could be tentatively related to CHC biosynthetic pathways. For *F. varia* and/or *C. analis*, functional terms related to pigmentation pathways (pigmentation, pigment metabolic process, pigment biosynthetic process, pigmentation during development, and terms related to eye pigments), were also significantly more represented in the Ne and Fg phases. These GO results (Fig 2) evidenced the similarities and differences in terms of cuticle-related functional attributes between the developmental phases and bee species. Some functional categories were shared by the three bee species, and a larger number of categories were shared by the two eusocial species than by one of them and *C. analis*.

**Fig 2.**
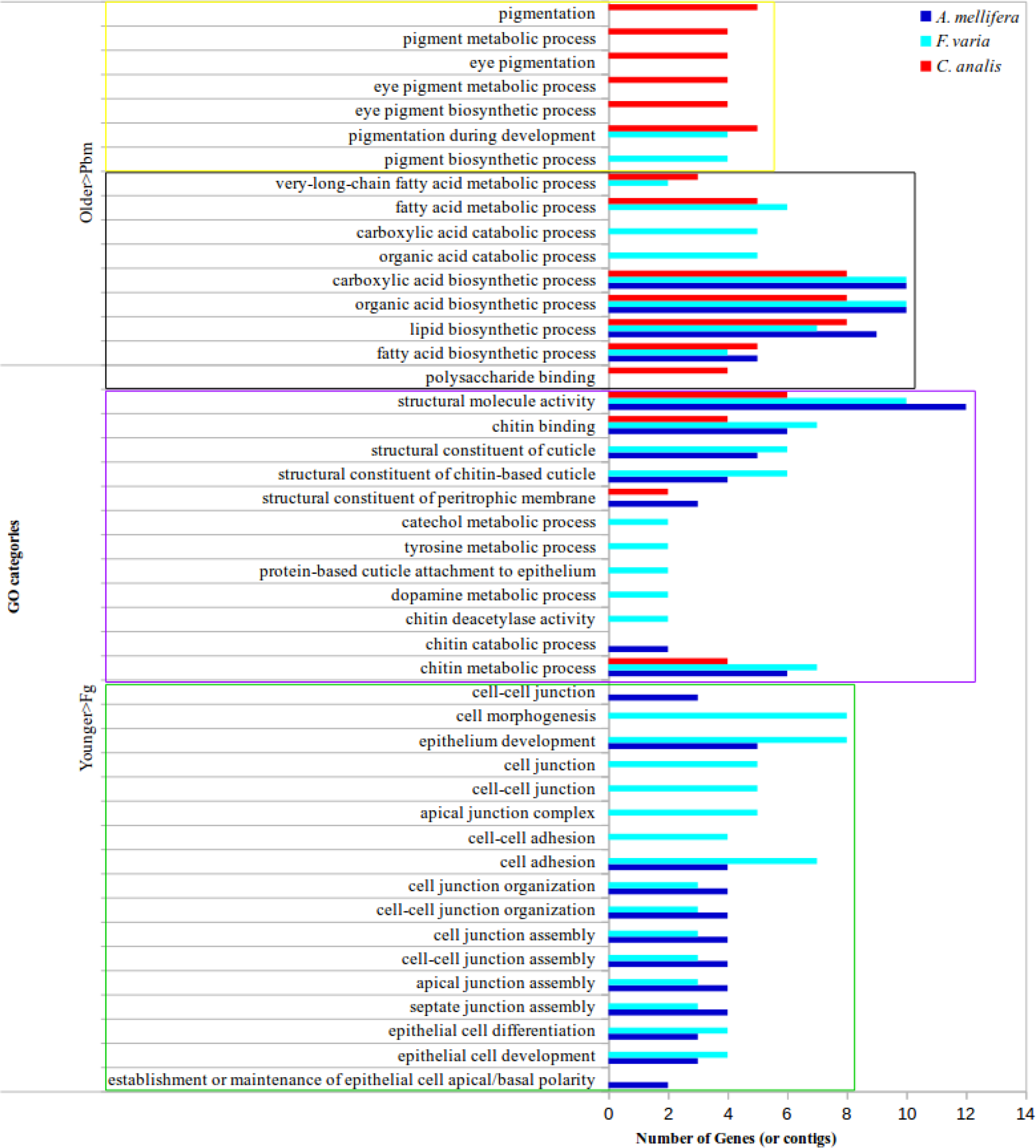
Gene Ontology (GO) functional terms attributed to integument genes during adult cuticle development and maturation. The functional terms more represented in the Pbm and Ne phases than in the Fg phase are indicated as Younger>Fg, and those more represented in the Ne and Fg phases than in the Pbm phase are reported as Older>Pbm. The green box includes GO terms related to the cuticle-producing tissue, the epidermis. Purple box: GO terms associated to structural components of the cuticle. Black box: GO terms potentially associated to CHC biosynthetic pathways. Yellow box: GO terms related to pigments and pigmentation.

S2 File specifies the genes upregulated between the developmental phases and bee species. Among the DEGs and DECs, it was clear that those encoding structural cuticular proteins, such as those in the CPR, Twdl, and Cpap families, and also chitin-related genes with roles in chitin metabolism, modification and degradation, were upregulated in the Pbm and/or Ne phases of the three bee species here studied. A series of sequences containing the chitin-binding peritrophin A domain were similarly overrepresented in the integument of the Pbm and/or Ne phases of *F. varia* and *C. analis*, thus being candidates to participate as structural proteins or enzymes in cuticle formation. In contrast, genes encoding odorant-binding proteins that bind to pheromones thus serving as insect chemoreceptors, as well as genes encoding a variety of CYPs (cytochrome P450), and antioxidant proteins like glutathione-S-transferase (GST), glutathione peroxidase (GTPx), thioredoxin peroxidase (TPX), and superoxide dismutase (SOD), were more expressed in the mature integument of foragers of the three bee species. Transcripts for genes related to the activity of juvenile hormone (JH), which is produced in a greater quantity in foragers [41], specifically *Krüppel homolog 1* (*Kr-h1*), and *JH-esterase* (*jhe*), were found in higher levels in the Fg integument of *F. varia* and *C. analis* than in the younger phases; transcripts for a JH-inducible (JHI-1) protein were overrepresented in the Fg and Ne integuments of *A. mellifera* in comparison to the Pbm integument. The Fg integument of *C. analis* showed a higher expression of the *ecdysone receptor* (*EcR*) and *seven-up*, an orphan nuclear receptor belonging to the steroid receptor gene superfamily [42]; *seven-up* is also overexpressed in the Fg integument of *F. varia*. Defense response genes (*defensin, apidaecin*) were also highly expressed in the Fg integument of *A. mellifera* (S2 File). Such developmental differences in gene expression in the integument reflect the dynamics of cuticle formation and acquisition of its functionality in adult bees

### Distance correlation analysis based on the RNA-seq data is consistent with the earlier cuticle maturation in the solitary *C. analis* in comparison to the eusocial *A. mellifera*

We used the DEGs and DECs in a distance correlation analysis in order to measure the clustering potential of the studied developmental phases of each bee species (Fig 3). This strategy allowed us to know for each of the bee species how near, or distant from each other are the Pbm, Ne and Fg developmental phases in terms of gene expression levels/patterns in the integument. Assuming that the cuticle of solitary bee species is sufficiently mature at the emergence, the hypothesis approached here was that the integument samples of the Ne and Fg phases of *C. analis* would cluster together, and separately from the Pbm samples. In contrast, in the eusocial species, the Pbm and Ne samples would group together, with the Fg samples forming a more distant group. Indeed, the results of the distance correlation analysis using all the *C. analis* DECs and *A. mellifera* DEGs were consistent with this hypothesis. In terms of differential gene expression in the integument, the Ne and Fg phases are nearest to each other in the solitary bee than they are in the eusocial *A. mellifera*. However, in *F. varia*, the distance correlation analysis grouped the Ne and Fg phases in a statistically supported cluster, in spite of the very distinct cuticle melanization patterns and hardness that they exhibit.

**Fig 3.**
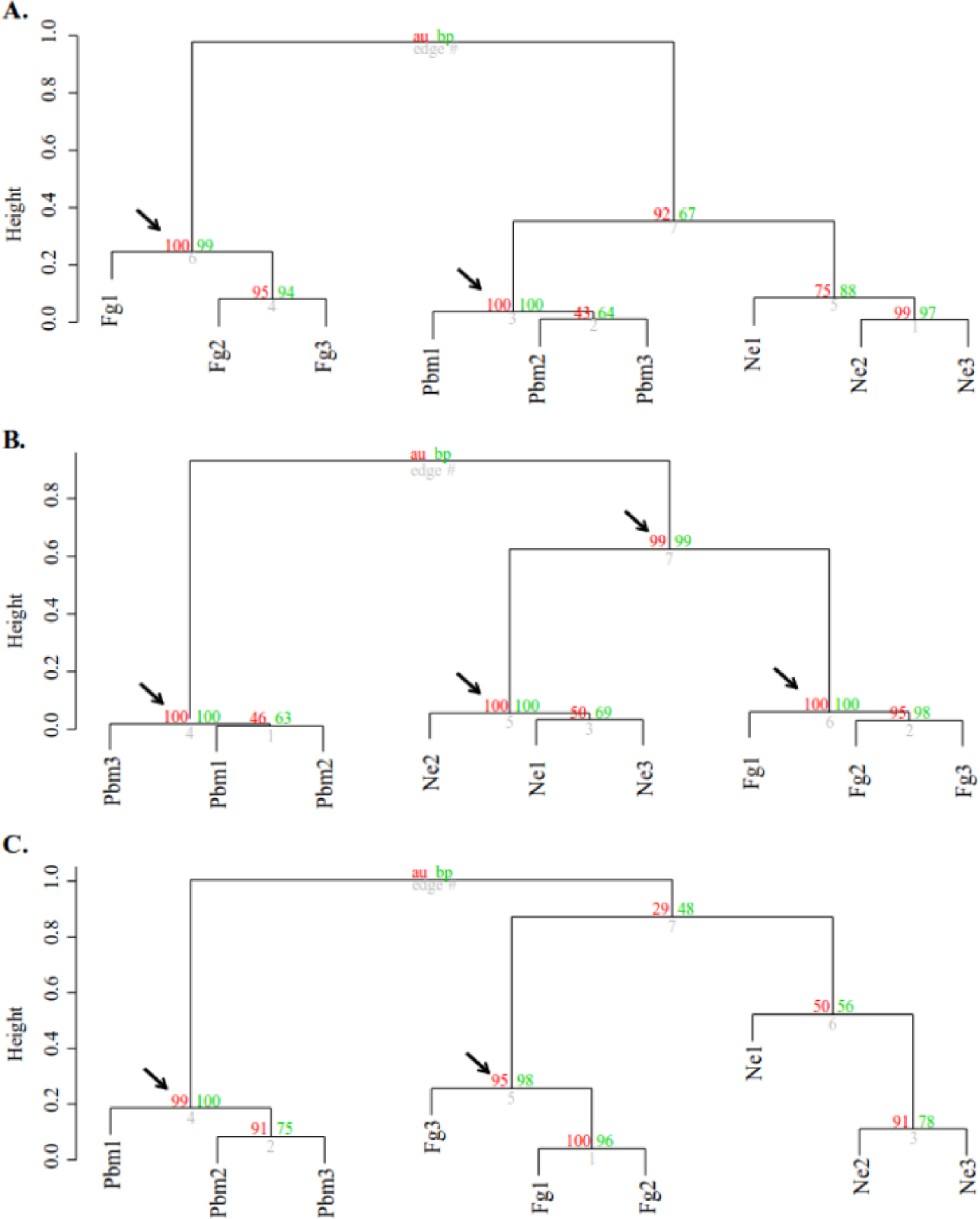
Distance correlation analysis between developmental phases based on the expression of DEGs and DECs. (A) *A. mellifera*; (B) *F. varia*; (C) *C. analis*. Red values (BP): bootstrap support. Green values (AU): cluster support. Arrows point to significant clusters (AU > 95%). Branch edges are shown in gray. Pbm = pharate adults. Ne = newly emerged bees. Fg = foragers.

### Gene expression profiles in the integument of the eusocial (*A. mellifera* and *F. varia*) and solitary (*C. analis*) bee species

Heatmaps representing the expression profiles of classes of cuticle-related genes through the Pbm, Ne and Fg developmental phases were constructed and clearly showed differences between the bee species (Fig 4).

**Fig 4.**
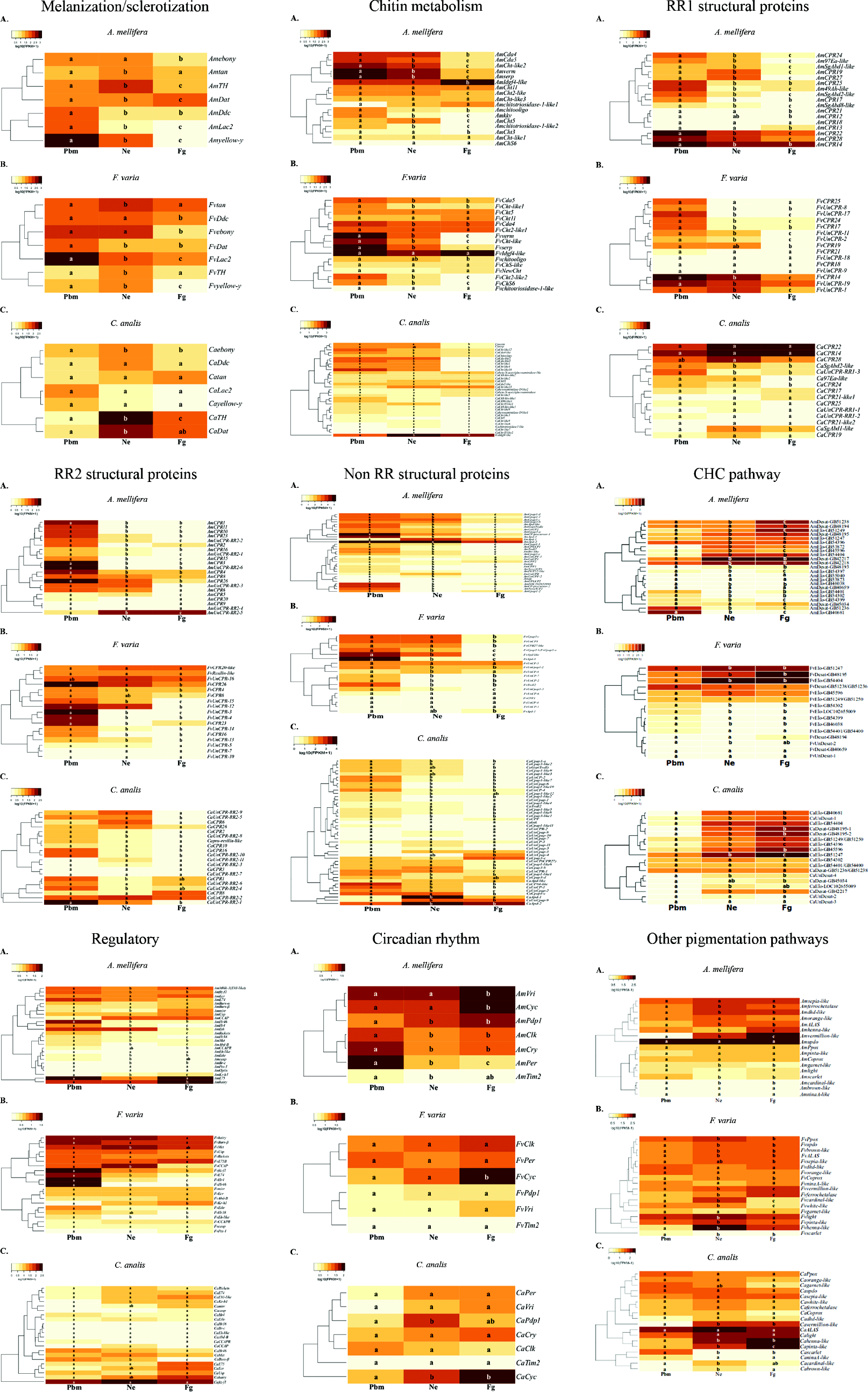
Representative heatmaps of gene expression profiles through the Pbm (pharate-adult), Ne (newly-emerged), and Fg (forager) developmental phases of (A) *A. mellifera*, (B) *F. varia*, and (C) *C. analis*. Genes were grouped according to their potential function in adult cuticle formation and maturation. Different lowercase letters on the heatmaps means statistically significant difference (see Materials and Methods) in the expression levels between the developmental phases of each bee species.

We found in the RNA-seq libraries seven genes involved in the biosynthesis of melanin and sclerotizing compounds (see a representation of the melanin/sclerotin biosynthetic pathway in Shamim *et al.* [24]. The genes with roles in the melanization/sclerotization pathway, except for *Dat*, were more expressed in the younger phases (Pbm and/or Ne) of *A. mellifera*. Similarly, these genes, including *Dat*, were more expressed in the younger phases of *F. varia*. In contrast, in *C. analis*, the majority of the genes in this class (*tan*, *Ddc*, *Lac2*, *yellow-y*) did not significantly change their expression levels, *ebony* was highly expressed in the Ne and Fg phases, and the expression profile of *Dat* also differed from both eusocial species*. TH* was the only gene in this class showing a significantly higher expression level in the very same developmental phase (Ne) of the three bee species.

Searching for genes related to pigmentation pathways other than the melanin biosynthetic pathway in the integument RNA-seq libraries, such as those genes involved in pterin, ommochromes, and heme formation, we found 17 genes in *A. mellifera*, and 18 genes in *F. varia* and also in *C. analis*, including *cardinal*, *scarlet, brown*, *vermillion*, *light*, *sepia*, and *henna* (this one involved in both biopterin formation, and tyrosine formation for the melanization process), thus indicating that their products are necessary in the adult cuticle. We also observed that a higher proportion (66.7%) of these genes displayed higher expression levels in the adults (Ne, Fg, or both phases) of *F. varia* in comparison to *A. mellifera* (29.4%) and *C. analis* (27.8%).

Concerning genes encoding chitin-related enzymes, we found 17, 16 and 33 of these genes in *A. mellifera*, *F. varia* and *C. analis*, respectively. The four *Cda* genes (*Cda4*, *Cda5*, *verm* and *serp*) found in the eusocial species, and five (*Cda4-like*, *Cda-like-1*, *Cda-like-2*, *verm*, *serp*) of the six *Cda* genes found in *C. analis* showed the higher expression in the Pbm, Ne, or both developmental phases, the other *C. analis Cda* gene (*Cda5-like*) showed no significant expression levels variation throughout the developmental phases. One of the two *ChS* genes found in *A. mellifera* (*kkv*) and *F. varia* (*ChS6*), and two of the four *ChS* genes found in *C. analis* (*ChS-kkv-like-1*, *ChS6-like1*) were also more expressed in the Pbm and/or Ne phases whereas the other *ChS* genes of the three bee species did not show significant expression level variation. Six (*Cht-like2*, *Cht-2 like*, *chitooligosaccharidolytic-domain-like*, *Cht5*, *chitotriosidase*, *Cht3*) of the eleven *Cht* genes of *A. mellifera*, and four (*Cht-like1*, *Cht-like*, *chitooligosaccharidolytic-domain-like*, *Cht2-like2*) of the ten *Cht* genes of *F. varia* were also more expressed in the Pbm and/or Ne phases, the remaining showing no significant variation in expression levels, except for the chitinase-encoding gene, *Idgf-4*, which is significantly more expressed in *A. mellifera* foragers. In contrast, only a small number (*Cht-like12*, *Cht-like4*, *Cht-like10*, *Cht-like1*) of the 22 *Cht* genes of *C. analis* were more expressed in these phases, the remaining showing no significant changes in expression levels, except for *Idgf-4*, which is more expressed in foragers.

The majority of the CPR genes (encoding cuticle proteins containing the RR1 or RR2 Consensus types) in the eusocial species showed significant variation in expression levels through the studied developmental phases, the proportions of RR1 and RR2 genes showing variable expression corresponding to 94.1% and 80.9% in *A. mellifera*, and 66.7% and 70.6% in *F. varia*, respectively. In contrast, lower proportions of RR1 and RR2 genes in *C. analis*, corresponding to 40% and 35% respectively, showed significant variation in transcript levels. For the three bee species, most of the genes showing changing transcript levels, in the range of 75 to 100%, were more expressed in the Pbm or both Pbm/Ne phases. Interestingly, a few CPR genes were significantly more expressed in the Ne phase (*AmCPR19*, *AmCPR27*, *AmSgAbd1-like*, *FvUnCPR-1*), or in both Ne and Fg phases (*CaSgAbd1-like* and *AmUnCPR-RR2-5*), and only a CPR gene, the RR1 motif *AmCPR13* gene, showed a higher expression exclusively in foragers. Similarly, a higher proportion of the non-RR cuticular protein genes showed significant transcript levels variation in *A. mellifera* and *F. varia*, 90.6% and 72.2% respectively, in comparison to *C. analis* (64.5%). These genes were also mostly more expressed in the Pbm or Pbm/Ne phases of the three bee species. However, like some CPR genes, there were non-RR genes displaying the highest expression in adults (Ne, Fg or both phases), specifically, *Apd* genes in *F. varia* (*FvApd-1*) and *C. analis* (*CaApd-1* and *CaApd-2*), and *Cpap* genes in *C. analis* (*CaUnCpap-3*, *CaUnCpap-4*, *CaUnCpap-9*, *CaCpap3-e*).

For the three studied bee species, a higher proportion of genes encoding elongases (Elo-genes) and desaturases (Desat-genes) putatively involved in CHC biosynthesis were more expressed in adults (Ne, Fg, or both phases) than in the Pbm phase. However, in *C. analis*, a higher proportion (66.7%) of these genes increased significantly their expression levels from the Pbm to the Ne phase in comparison to *F. varia* (26.7%) and *A. mellifera* (39.1%).

A higher proportion of the regulatory genes was significantly more expressed in the Pbm phase of *A. mellifera* (50%) and *F. varia* (28.6%) than in *C. analis* (4.5%) in which the majority of the genes (72.7%) did not show significant difference in expression levels between the developmental phases. Some regulatory genes had a higher expression in adults (Ne, Fg or both phases) of *A. mellifera* [*Ammirr* (*mirror*), *AmUsp* (*Ultraspiracle*)*, AmCCAP* (*Crustacean Cardioactive Peptide*), *AmKr-h1* and *Amhairy*], *F. varia* (*FvKr-h1*), and *C. analis* (*CaKr-h1*, *Camirr*, *CaE75*, *CaEcR* and *Cahairy*). Two of the regulatory genes in *A. mellifera*, *AmE75* and *AmMblk* (*Mushroom body large type Kenyon cell specific protein-1* or *E93-like*), which were highly expressed in the younger Pbm phase, were also highly expressed in the older Fg phase.

Four among the seven circadian rhythm genes of *A. mellifera* [*Clk* (*Clock*), *Cry* (*Cryptochrome*), *Per* (*Period*) and *Tim2* (*Timeless2*)] showed the highest expression in the Pbm phase. This is in contrast to the majority of the circadian rhythm genes in *F. varia* [*Clk*, *Per*, *Pdp1* (*Par domain protein I*), *Vri* (*vrille*), *Tim2*] and *C. analis* (*Per*, *Vri*, *Cry*, *Clk*, *Tim2*), which did not significantly change their expression levels. The genes *Vri*, *Cyc* (cycle), and *Pdp1* in *A. mellifera*, *Cyc* in *F. varia*, and *Cyc* and *Pdp1* in *C. analis* showed the highest expression in adults (Ne, Fg or both phases).

In summary, the main differences between the social and solitary bee species were highlighted in the heatmaps (Fig 4) displaying integument gene expression profiles: **(a)**A higher proportion of genes involved in the melanization/sclerotization pathway, cuticle formation (RR1, RR2, and non-RR genes), and regulation (regulatory genes) showed significant transcript levels variation through the studied developmental phases of *A. mellifera* and *F. varia* in comparison with *C. analis*. Most of these genes showing transcript levels variation were more expressed in the Pbm or Pbm/Ne phases. In *C. analis*, the higher proportion of genes displaying no differences in expression levels through the studied phases, were possibly highly expressed earlier, before the Pbm phase, for faster cuticle formation and maturation, but this assumption requires further investigations; **(b)**The number of chitin-related genes, higher in *C. analis*, and not their expression patterns, distinguished this species from the eusocial *A. mellifera* and *F. varia*; **(c)**A higher proportion of desaturase and elongase genes putatively involved in CHC biosynthesis showed significantly increased expression levels at the emergence (Ne phase) of *C. analis* in comparison to the eusocial ones, which is consistent with an accelerated process of cuticle maturation in the solitary bee.

Importantly, all the gene classes here studied included representatives showing increased or high expression levels in the mature integument of foragers indicating that the mature cuticle is a dynamic structure requiring structural and regulatory elements for its maintenance.

### Correlation among expression profiles of genes candidates to play roles in cuticle formation/maturation in the eusocial (*A. mellifera* and *F. varia*) and solitary (*C. analis*) bees

We used Pearson’s correlation in order to measure the strength of the linear association between the expression profiles of 119 genes related to cuticle development and maturation shown in Fig 4, which shared potential orthology relationships between the bee species. A fraction of these ortholog genes showed non-significantly correlated transcript levels fluctuation among the bee species, thus highlighting peculiarities in cuticle development for each species. However, 76 orthologs (S1 Table; Fig 5) displayed expression profiles significantly correlated at least between two of the three bee species. Importantly, the expression profiles of 21 among these 76 genes were positively correlated between the eusocial species, and negatively or non-correlated with the solitary bee (r≥0.6 and p≤0.1). In addition, other six genes, whose transcripts were not identified in *C. analis*, showed expression profiles positively correlated between the eusocial species. Therefore, these 27 genes are possibly contributing to differences in the processes of cuticle development and maturation in the eusocial bees versus the solitary bee. Thus, the expression profiles of genes related to the melanization/sclerotization pathway (*ebony*, *tan*) and chitin metabolism [*Idgf4-like, Cda5* (*Chitin deacetylase 5*), *chitooligosacchariodolytic-domain-like*], genes encoding cuticular structural proteins containing the RR1 or RR2 domains (*CPR14*, *CPR17*, *CPR18*, *CPR23*, *CPR25*, *CPR26*), or lacking these domains (*Apd-3*, *Apd-like*), and also genes in CHC pathways (*Desat-GB40659*, *Elo-GB54401*, *Elo-GB54302*, *Elo-GB45596*, *Elo-GB46038*), regulators of cuticle development [*Ethr* (*Ecdysis triggering hormone receptor*), *E74*, *Hr4* (*Hormone receptor 4*), *Hr38* (*Hormone receptor 38*), *FTZ-F1* (*Fushi tarazu-factor 1*), *rickets*, *Ptx-1* (bicoid-related *Paired-type homeobox gene D*), circadian rhythm genes (*Tim2*) and a gene in the non-melanin pigmentation pathways, *ALA*S (*δ-aminolevulinic acid synthase*)], suggest roles in the differential cuticle development in the solitary versus eusocial bees.

**Fig 5.**
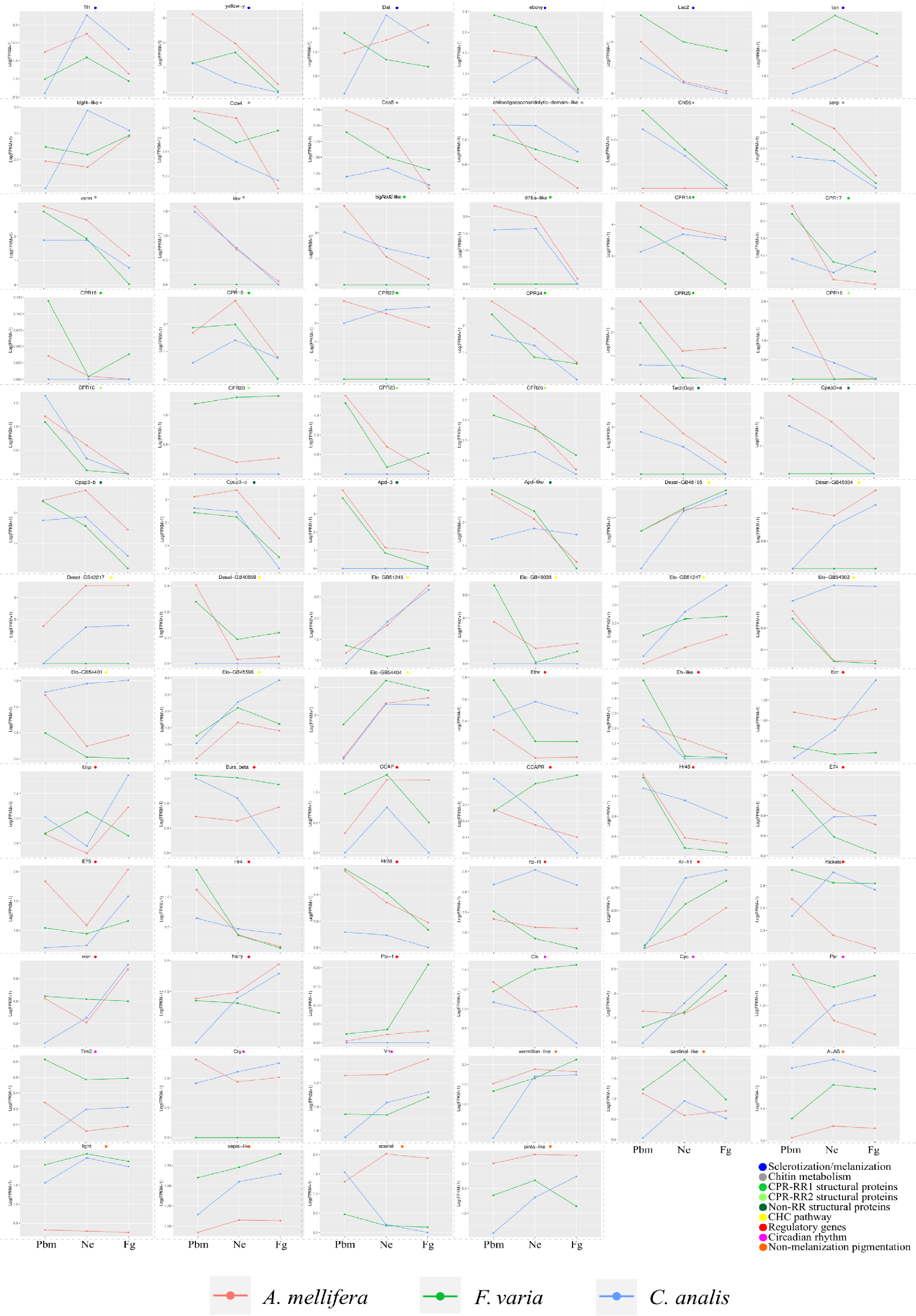
Ortholog genes showing significantly correlated expression profiles at least between two of the three bee species, *A. mellifera*, *F. varia* and *C. analis*. Expression profiles of *ebony*, *tan*, *Idgf4-like*, *Cda5*, *chitooligosaccharidolytic-domain-like*, *CPR14*, *CPR17*, *CPR25*, *CPR26*, *Apd-like*, *Elo-GB54302*, *Elo-GB54401*, *Elo-GB45596*, *Ethr*, *E74*, *Hr4*, *Hr38*, *FTZ-F1*, *rickets*, *Tim2*, and *ALAS* were positively correlated between the eusocial bee species, and negatively or non-correlated with the solitary bee. Expression profiles of *CPR18*, *CPR23*, *Apd-3*, *Desat-GB40659*, *Elo-GB46038*, and *Ptx-1* were positively correlated between the eusocial species, the basal line in the graphic representations indicating undetected orthologs in *C. analis*. Pbm: pharate adults, Ne: newly emerged, and Fg: foragers. Color key at the bottom of figure.

Among the above cited 76 orthologs, we also found genes whose expression profiles were positively correlated between the solitary and eusocial bees. Thus, the following 23 genes shared expression profiles positively correlated between *A. mellifera* and *C. analis*: *yellow-y* (melanization /sclerotization pathway), *Cda4* and *ChS-kkv-like1* (chitin metabolism), *SgAbd2-like* and *97Ea-like* (CPR-RR1 class), *CPR10* (RR2 class), *Twdl(Grp)*, *Cpap3-a*, *Cpap3-b* and *Cpap3-c* (non-RR class), *Desat-GB48195*, *Desat-GB45034*, *Desat-GB42217*, *Elo-GB51249* and *Elo-GB54404* (CHC pathways), *Usp*, *CCAPR*, *Mirr* and *hairy* (regulatory genes), *Cyc* (Circadian rhythm), *verm*, *sepia* and *pinta-like* (other pigmentation biosynthetic pathways than melanin). Similarly, the following 12 genes shared expression profiles positively correlated between *F. varia* and *C. analis*: *Cda4* and *ChS6* (chitin metabolism), *Cpap3-c* (non-RR class), *Elo-GB54404* (CHC pathways), *Bursβ*, *CCAP* and *E75* (regulatory genes), *Cyc* (Circadian rhythm), *verm-like*, *cardinal-like*, *light* and *scarlet* (other pigmentation biosynthetic pathways than melanin).

### Co-expression networks reconstructed with genes related to cuticle development and maturation, and common interactions between the networks of the eusocial *A. mellifera* and *F. varia* bees

The genes related to cuticle formation and maturation in *A. mellifera*, *F. varia*, and *C. analis* were separately used for co-expression networks reconstruction (S2-S4 Figs). The gene co-expression networks for the eusocial species, *A. mellifera* and *F. varia*, showed common interactions among regulatory elements [*FTZ-F1*, *E74*, *Hr4*, *Hr46* (*Hormone receptor 46*)], genes encoding structural cuticular proteins (CPR14, CPR17, CPR23, CPR24, CPR25, Apd-3 and Apd-like), and encoding the elongase Elo-GB54302, Cdas [*verm* (*vermiform*), *serp* (*serpentine*), *Cda5*], and Lac2 (Fig 6). However, by intersecting the gene co-expression networks of the eusocial *A. mellifera* and the solitary *C. analis*, we found only one common interaction comprising the genes *yellow-y* and *Cpap3-a*. Similarly, only the interactions between *CPR16* and *Eh-like* (*Eclosion hormone*), and *tan*/Elo-GB45596 were highlighted as being common to the eusocial *F. varia* and the solitary *C. analis* after superimposing their respective gene co-expression networks.

**Fig 6.**
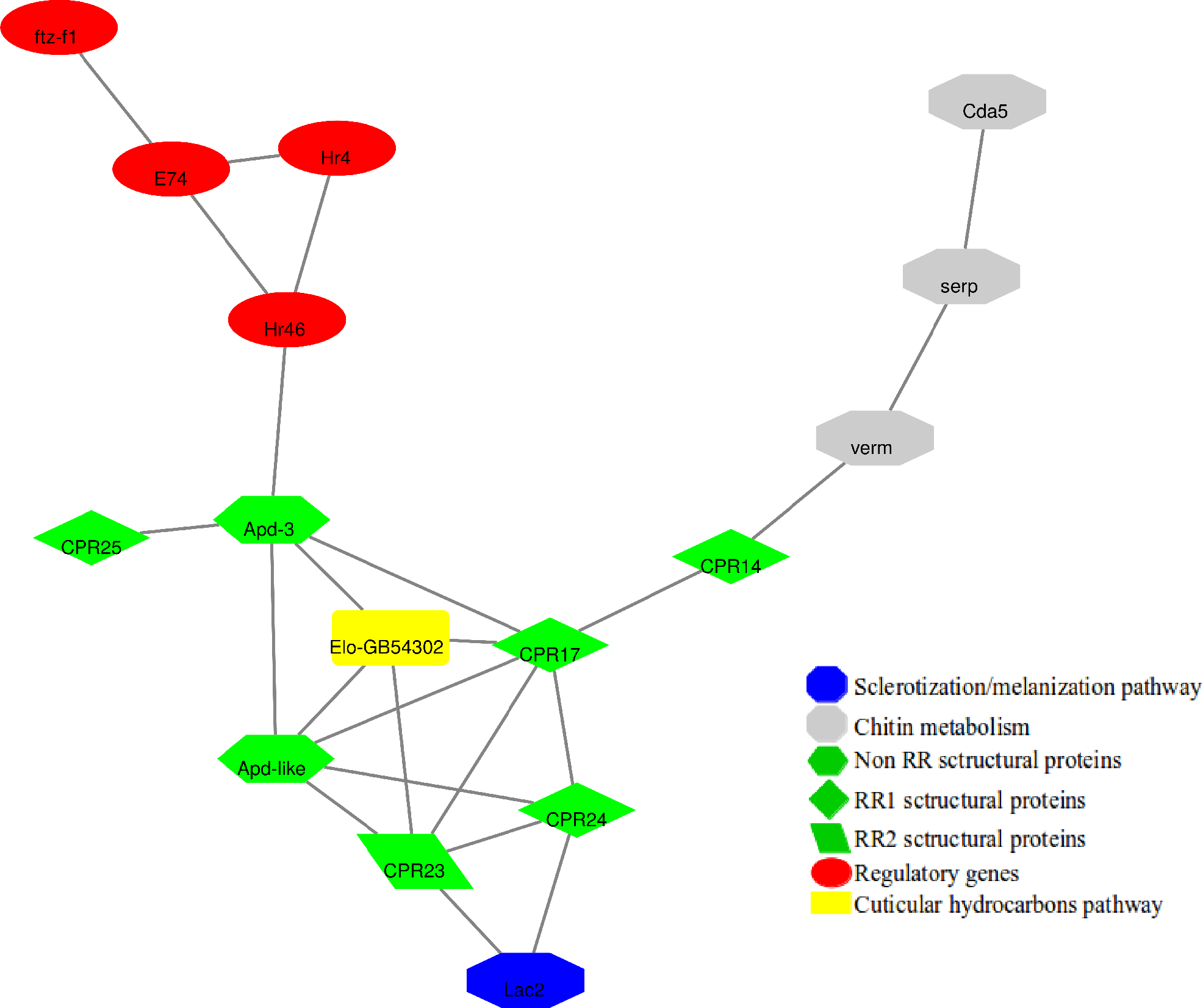
Overlapping interactions in the gene co-expression networks reconstructed with *A. mellifera* (S2 Fig) and *F. varia* (S3 Fig) genes related to cuticle formation and maturation.

### Ultrastructure and thickness of the developing adult cuticle shows conspicuous differences among the eusocial, primitively eusocial, facultatively eusocial, and solitary bee models

The morphology of the developing adult cuticle is shown for the eusocial *A. mellifera* and *F. varia* bees, for the primitively eusocial *Bombus brasiliensis*, for the facultatively social *Euglossa cordata*, and for two solitary bees, *C. analis* and *T. diversipes* (Fig 7). For *A. mellifera*, there were no noticeable modifications in cuticle ultrastructure from the pharate-adult phase (Pbm) to 48h after emergence. Up to this time, only the exocuticle was deposited. At 72h, endocuticle layers became apparent in the micrographs (Fig 7A). Cuticle ultrastructure was very similar in 96h-aged *A. mellifera* bees and foragers (Fig 7A). We then measured the thickness of the cuticle in seven time points of *A. mellifera* development (Fig 7A’). As the cuticle measurements in the groups of bees aging 0h to 96h post-emergence, and in the group of foragers, did not show a normal distribution (Shapiro-Wilk normality test, p = 0.0074) we used the Kruskal-Wallis test associated with the *post hoc* Conover-Iman test and Bonferroni correction to compare the sample collection data. Foragers have a significantly thicker cuticle in comparison to the earlier developmental phases, i.e., the Pbm phase, and bees at 0h, 24h and 48h after emergence (Fig 7A’). At 72h and 96h post-emergence, cuticle measurements values did not significantly differ from foragers. Differently, the cuticle of the eusocial *F. varia* showed very little variation in morphology (Fig 7B), and no significant variation in thickness (Fig 7B’) from the Pbm phase to the forager time. For the solitary species, *C. analis*, we observed remarkable differences in cuticle ultrastructure (Fig 7C) and thickness (Fig 7C’) between the Pbm and Ne phases, whereas the cuticles of the Ne and Fg phases were very similar. Pore canals are abundant in the Pbm cuticle of *C. analis*. At the Ne and Fg phases, the *C. analis* cuticle can be described as a succession of lamellae, the most superficial ones, i.e., those first deposited, became thicker and reached a higher degree of differentiation (Fig 7C). Like *C. analis*, the cuticle of *B. brasiliensis* (Fig 7D, 7D’), *E. cordata* (Fig 7E, 7E’), and *T. diversipes* (Fig 7F, 7F’), did not show noticeable ultrastructural changes, or statistically significant thickness differences, from the emergence (Ne phase) to the forager time (Fg phase).

**Fig 7.**
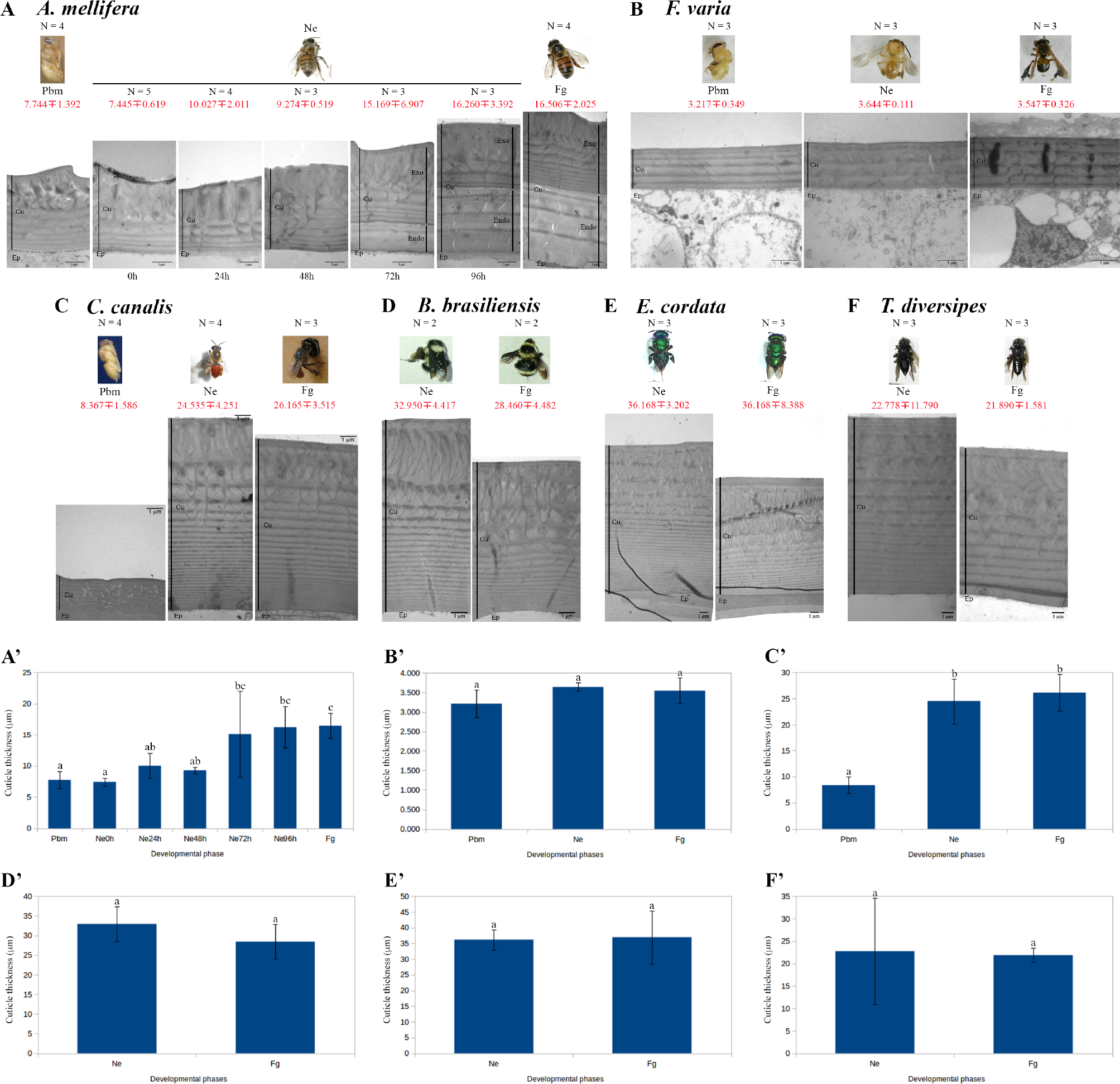
Ultrastructure and thickness of the developing and mature adult cuticle of bees differing in ways of life. (A) *A. mellifera* (eusocial), (B) *F. varia* (eusocial), (C) *C. analis* (solitary), (D) *B. brasiliensis* (primitively eusocial), (E) *E. cordata* (facultatively eusocial), and (F) *T. diversipes* (solitary). Developmental phases are indicated: Pbm (pharate-adult); Ne (newly-emerged); 0h, 24h, 48h, 72h, and 96h after adult emergence; Fg (forager). The cuticle/epidermis junction was used to align the cuticle images. Means and standard deviations of cuticle thickness measurements are represented in red. The number of cuticle samples measured (N) is indicated for the Pbm, Ne and Fg phases of each bee species. (A’-F’) Cuticle thickness measurements (µm) for the corresponding bee species. Different lowercase letters indicate significant statistical difference between the developmental phases of each species.

Together, these data indicate that cuticle deposition in the solitary species, *C. analis* and *T. diversipes*, and the primitively and facultatively eusocial species, *B. brasiliensis* and *E. cordata*, respectively, is completed or almost completed at the time of adult emergence. In contrast, in *A. mellifera*, the endocuticle was deposited only after the emergence. Surprisingly, the cuticle of the eusocial *F. varia* species did not undergo significant variation in ultrastructure and thickness from the Pbm to the Fg phases, although a great increase in pigmentation and sclerotization has been clearly noticed in *in vivo* observations.

### Cuticular n-alkanes mark the earlier cuticle maturation in the solitary *C. analis* compared to the eusocial *A. mellifera* and *F. varia* bee species

The CHC composition of the superficial cuticle layer, the envelope, was determined for *A. mellifera*, *F. varia* and *C. analis* as another strategy potentially able to uncover differences that could be associated to the cuticle maturation heterochrony. The proportion of CHCs in the chromatograms, the significance level of each peak and the contribution of these peaks for discriminating the developmental phases of the eusocial and solitary species are shown in S4 File. The Euclidean distance clustering analysis applied to the total CHC quantification data clearly discriminated the Fg phase from the earlier Pbm and Ne phases in the eusocial bees, *A. mellifera* and *F. varia*, as well as in the solitary *C. analis* (S5 Fig). Total CHC quantification data grouped together the Pbm and Ne samples of *A. mellifera* (AU=100; BT=100), *F. varia* (AU= 100; BT=100), and *C. analis* (AU=96; BT=88). For *F. varia*, the group including Ne samples showed AU=95 and BT=87, which is a moderate to high BT value usually associated with Bayesian posterior probabilities ≥ 95% [43]. The same was verified for the *F. varia* Pbm samples (AU=94; BT=87). For the two eusocial species, the Fg samples grouped with maximal AU (100) and BT (100) values. For *C. analis*, however, these values were significantly lower (AU=78; BT=57) (S5 Fig).

When we analyzed separately the CHC classes, n-alkanes discriminated the *A. mellifera* foragers (Fg) (AU=94; BT=84) from the Pbm and Ne developmental phases, which were clustered together (AU=85; BT=77). As in *A. mellifera*, n-alkanes also discriminated the *F. varia* foragers (Fg) as a separate group (AU=97; BT=76), and the Ne and Pbm phases were clustered together (AU=98; BT=77). However, the n-alkanes did not significantly distinguish the developmental phases of *C. analis* (S5 Fig).

The unsaturated CHCs data from *A. mellifera* did not give us back a strong support for distinguishing the developmental phases. Although all the Ne samples and the majority of the Pbm samples have been grouped with a high AU value (99%), the BT=1 value was low. Three of the *A. mellifera* foragers (Fg) escaped from the main cluster formed by twelve foragers (AU=96; BT=23). In contrast, the unsaturated CHCs discriminated each of the developmental phases of *F. varia*. The groups of Pbm samples (AU=99; BT=93) and Ne samples (AU=99; BT=94) were maintained together in a larger cluster (AU=99; BT=98), and separately from the group of Fg samples (AU=99; BT=97). This CHC class clustered together the Pbm and Ne samples of *C. analis* (AU=96; BT=80). The Fg samples of *C. analis* were separated into two main clusters, respectively supported by AU=94; BT=70 and AU=93, BT=83 (S5 Fig).

Branched CHCs from *A. mellifera* clearly clustered the Fg samples (AU=97; BT=94). The Ne and Pbm phases were joined together in a single well-supported group (AU=100; BT=100). In *F. varia*, separation of Fg from the earlier phases was not clear: three of the fifteen Fg samples joined to the group encompassing the Pbm and Ne samples, this group being supported by 98% AU, but showing a low BT value (BT=3). The *F. varia* forager samples were also clustered with low BT values. In the solitary *C. analis*, the branched CHCs clustered six of the seven Fg samples into a single group (AU= 97; BT=72), and all the Ne samples plus two of the four Pbm samples were clustered together in another group supported by AU=99, but presenting a low BT value (BT=39) (S5 Fig).

These data on the Euclidean distance based on the relative quantification of CHCs was contrasted with the results on the absolute quantification of CHCs (CHC µg per bee) (Table 1; S4 File). Table 1 shows that Fg bees of the eusocial species have significantly higher quantities of n-alkanes than the Ne and Pbm bees, which is not true for *C. analis*. In addition, absolute quantification of unsaturated CHCs also distinguished the foragers from the earlier developmental phases of *A. mellifera*, but not of *C. analis*. For *F. varia*, the mass of unsaturated compounds could not be quantified due to their very low quantities.

**Table 1.** Absolute quantification of n-alkanes and unsaturated CHCs in the cuticle of eusocial and solitary bee species. Developmental phases are indicated: Pbm (pharate-adults), Ne (newly emeged bees), Fg (foragers). Means and standard deviations (STD) of 3 samples (N=3) per developmental phase. Different lowercase letters in the Sig (statistical significance) column indicate difference between the developmental phases of each species.

| N-alkanes |  |  |  |  |
| --- | --- | --- | --- | --- |
| *A. mellifera* |  |  |  |  |
| Phase | Mean | ± | STD | Sig. |
| Pbm | 9.35965938 | ± | 2.89275421 | **a** |
| Ne | 7.36272669 | ± | 1.44535498 | **a** |
| Fg | 18.2360314 | ± | 4.35877417 | **b** |
| *F. varia* |  |  |  |  |
| Phase | Mean | ± | STD | Sig. |
| Pbm | 1.63324436 | ± | 0.15390427 | **a** |
| Ne | 3.40142407 | ± | 1.35387231 | **a** |
| Fg | 9.28337077 | ± | 3.03839358 | **b** |
| *C. analis* |  |  |  |  |
| Phase | Mean | ± | STD | Sig. |
| Pbm | 6.56549413 | ± | 1.62457012 | **a** |
| Ne | 14.8349947 | ± | 0.32610609 | **b** |
| Fg | 13.3273848 | ± | 5.07830924 | **ab** |
| Unsaturated CHC |  |  |  |  |
| *A. mellifera* |  |  |  |  |
| Phase | Mean | ± | STD | Sig. |
| Pbm | 0.60042039 | ± | 0.17210242 | **a** |
| Ne | 0.92421769 | ± | 0.09047864 | **a** |
| Fg | 6.5543118 | ± | 2.38207067 | **b** |
| *C. analis* |  |  |  |  |
| Phase | Mean | ± | STD | Sig. |
| Pbm | 9.24954719 | ± | 2.48578756 | **a** |
| Ne | 19.242 | ± | 2.40516304 | **ab** |
| Fg | 28.3380901 | ± | 11.855003 | **b** |

Standard deviation (STD). Different red letters in significance (Sig.) column represent statistical significance (ANOVA associated to Tukey's HSD post hoc test) between developmental phases of a species.

In summary, the Euclidean distance analysis based on the relative quantifications of n-alkanes, as well as the absolute quantifications of n-alkanes and unsaturated CHCs, were consistent with the hypothesis of interdependence between cuticle maturation timing and the eusocial/solitary ways of life. These analyses distinguished the foragers from the younger bees, but only in *A. mellifera* and *F. varia*, this being interpreted as the cuticle achieving its complete maturation tardily in the eusocial species, whereas the solitary bee emerges with an already mature cuticle.

## Discussion

The RNA-seq analysis revealed the set of genes expressed in the integument of three bee species, and also the changes in gene expression as the adult cuticle is deposited and differentiates in a mature and fully functional cuticle. For *A. mellifera*, for which we have the sequenced genome, the genes expressed in the integument represented 95.07% of the genes in the released genome assembly version 4.5. Similar proportions will likely be found for *F. varia* and *C. analis* in the near future, after the sequencing of their respective genomes. Selected genes with potential roles in cuticle formation and maturation were characterized in terms of differential expression profiles. Co-expression networks were reconstructed. In parallel, we examined the ultrastructure of the developing adult cuticle of bee species. Furthermore, the CHC composition of the envelope, the less known cuticle layer, was also characterized. Our data expanded the knowledge on the insect integument. It is our expectation that the obtained data provide a valuable resource for future studies on exoskeleton formation and maturation in arthropods.

### Expression profiles of cuticle-related genes may significantly differ during adult cuticle formation/maturation, and among bee species

Genes involved in adult cuticle formation in *A. mellifera* in general show higher expression soon after the ecdysteroid titer peak that signalizes pupal cuticle apolysis and the beginning of the pharate-adult stage [44, 45]. Consistently, the majority of the integument genes showing expression levels variation in the three bee species, and identified as playing roles in cuticle melanization/sclerotization, cuticle structure (RR1, RR2, and non-RR genes), and regulation of the molting events (regulatory genes), displayed a higher expression in pharate-adults (Pbm phase), sometimes extending their higher expression up to the emergence time (Ne phase). However, we found genes, including those related to melanization/sclerotization and other pigmentation pathways, and also genes related to chitin metabolism, and structural cuticle protein genes, which showed the highest expression later, at emergence (Ne phase), and even in foragers (Fg phase), suggesting that their products are incorporated into the mature cuticle. Moreover, all transcripts identified in higher quantities during cuticle formation in pharate-adults were also identified in the newly emerged and forager bees, although in lower quantities. Their products may be involved in adult cuticle maintenance. Our gene expression findings indicate that the structure of the mature cuticle entails a dynamism, which has been up to now mainly characterized in studies on CHC composition of its most superficial layer, the envelope [23, this work].

Among the genes identified in the RNA-seq analysis of the integument, we focused on classes of genes playing roles in cuticle formation and maturation, such as those below discriminated.

### Genes related to cuticle pigmentation and sclerotization

The expression patterns of the first gene in the pigmentation/sclerotization biosynthetic pathway, *TH*, were positively correlated between *A. mellifera*, *F. varia* and *C. analis*, and apparently, *TH* does not contribute to the differential timing of cuticle pigmentation among them. Lower levels of *TH* transcripts were verified for the forager bees of the three bee species, which is consistent with the reported reduction in *TH* transcripts levels in *T. castaneum* [46, 47] and *Diacamma* sp [48] following the emergence. However, the expression patterns of *ebony* and *tan*, whose protein products act in a reversible reaction between dopamine and NBAD sclerotin [49], were positively correlated exclusively between the eusocial species, thus differentiating these species from the solitary one. The expression profiles of the remaining genes in the melanization/sclerotization pathway, including the *Lac2* gene previously characterized in *A. mellifera* [50], did not show such correlation patterns. Interestingly, *Dat* showed significantly increased expression in the mature cuticle of *A. mellifera* foragers, which is an uncommon pattern for genes in the melanization/sclerotization pathway.

We also observed that in general, the genes involved in the biosynthesis of other pigments except melanin displayed a higher expression levels in adults (Ne, Fg, or both phases) of *F. varia*, which may be tentatively interpreted as these genes playing roles in the process of post-ecdysial cuticle pigmentation in this bee species. Two of these genes, *cardinal* and *scarlet*, are both necessary for ommochromes formation in *B. mori* [51], and are associated to the formation of red and brown pigments [52]. The expression profiles of *light*, which is required for pigment granules formation [53], were positively correlated in *F. varia* and *C. analis*, and might be related to the brownish and reddish color pattern typical of the cuticle of these two species. The expression profiles of the gene encoding *ALAS*, which catalyzes the first enzymatic step in heme biosynthesis, were positively correlated exclusively between the eusocial species, *F. varia* and *A. mellifera*. *ALAS* might be involved in detoxification, as suggested for *D. melanogaster* [54, 55], and in prevent dehydration [56]. Interestingly, in contrast to the eusocial bees, the expression of *ALAS* is higher in the Pbm phase of *C. analis*, which may suggests that mechanisms of protection against cuticle dehydration develop anticipatedly in the solitary species.

### Genes involved in chitin synthesis, modification and degradation

In insects, *Cht*, *Cda*, and *ChS* genes have been described as highly expressed during cuticle renewal at the pharate-adult development [8, 57–62]. This was also observed in the bee species here studied, but with variations: the expression of a putative chitinase, *Idgf4-like*, increased in newly-emerged *C. analis*, and like reported for *T. castaneum* [63], this may be important for the transition to the adult stage. In *A. mellifera* and *F. varia*, the expression of *Idgf4-like* is high in foragers, supporting roles in the mature adult cuticle. Therefore, the decay in the expression of chitinase genes in adult insects seems not a standard pattern. Concerning the *Cda* genes, in *Drosophila*, they have a strict relationship with the mechanical properties of the exoskeleton [64], and this might be true for the *Cda* genes expressed in the integument of the bee species. The other class of chitin-related genes encodes ChS enzymes, which catalyze the last step in the chitin biosynthetic pathway and have been implied in the synthesis of epidermal cuticle in *T. castaneum* [65]. A *ChS* gene, *CS-1*, also called *krotzkopf verkehrt* (*kkv*), is required for procuticle formation, stabilization of the epicuticle, and attachment of the cuticle to the epidermis in *D. melanogaster* [66]. We found a *kkv* gene in *A. mellifera* (*Amkkv*) and three potential orthologs in *C. analis* (*CaChS-kkv-like 1*, *CaChS-kkv-like 2*, *CaChS-kkv-like 3*); this gene was not identified in the *F. varia* integument transcriptome.

### Genes encoding structural cuticular proteins

The large number of different cuticular protein genes found in insect genomes suggested that their products display redundant and complementary functions [67]. A variable number of genes encode the different classes of structural cuticular proteins in the three bee species and other hymenopterans (S2 Table). Thirty-two CPR genes had been previously identified in *A. mellifera* [12]. We detected other six CPR genes in our RNA-seq analysis of the *A. mellifera* integument, and also 32 and 35 CPR genes in the integument of *F. varia* and *C. analis*, respectively. In addition to have roles as structural proteins in the horizontally arrayed cuticular laminae, the function of some CPR proteins in *T. castaneum* was associated to the formation and organization of the pore canals vertically extended across the cuticle [68, 69]. This finding and the variety of CPR genes identified up to now suggest that distinct and additional functions are yet to be discovered for members of the CPR protein class.

Like the class of CPR proteins, Twdl proteins are structural cuticular components that effectively bind chitin, as demonstrated in *Bombyx mori* [70]. Two *Twdl* genes were previously characterized in the thoracic integument of *A. mellifera* [44], and now in the abdominal integument, thus indicating that Twdl proteins participate of both rigid (thoracic) and more flexible (abdominal) cuticles. Like *A. mellifera*, *C. analis* has two *Twdl* genes, but we identified only one in *F. varia*.

Two CPLCP-encoding genes as reported in Willis [12] were herein confirmed in *A. mellifera*. Genes in this family were identified in insect genomes in general and are very enriched in mosquito genomes [71]. Based on sequence homology, we could not identify *CPLCP* transcripts in the *F. varia* and *C. analis* abdominal integument.

CPF proteins were associated to the outer cuticle layers of *A. gambiae* and, apparently, do not bind chitin [72]. Three *CPF* genes were previously reported for *A. mellifera* [12] and one of them, *AmCPF1*, was validated in the thoracic integument through microarray analysis [45]. Here we found *CPF1* and *CPF2* transcripts in the abdominal integument of *A. mellifera*, and in addition, transcripts for two other CPF proteins, *AmUnCPF1* and *AmUnCPF2*. We also identified one *CPF* gene in *F. varia* and one in *C. analis*.

*Apd* genes seem exclusive of hymenopterans and three of these genes were previously identified in *A. mellifera* [73]. Their transcript levels in the thoracic integument were higher in pharate-adults compared to earlier developmental phases [45]. Here, we detected one more *Apd* gene in *A. mellifera*, *AmApd-like*, and three *Apd* genes in *F. varia* as well as in *C. analis*.

Cpap proteins are essential for the correct formation of the cuticular exoskeleton and elytra in *T. castaneum* [74]. In our RNA-seq analysis, we identified transcripts of three *Cpap1* genes (encoding Cpap proteins containing one chitin-binding domain) in *A. mellifera* and two *Cpap1* genes in *F. varia*, and also verified that the *C. analis* integument is very enriched in *Cpap1* transcripts (n=12), and also in *Cpap* transcripts (n=11) that we could not classify as encoding Cpap1 or Cpap3 (containing three chitin-binding domains). The number of *Cpap3* genes (5 genes) in *A. mellifera* [12] is here confirmed, and two and seven *Cpap3* genes were found in the *F. varia* and *C. analis* integument transcriptomes, respectively. It is important to observe that the genes originally named as *Am-C* and *Am-D* by Soares *et al.* [45] were here renamed as *AmCpap3-c* and *AmCpap3-d*.

The genes, *dumpy* (*dp*), *knk* (*knickkopf*) and *Rtv* (*Retroactive*) have also been identified as encoding cuticular proteins. In *D. melanogaster*, *dp* play roles in cuticle formation [12]. We detected transcripts for *dp* in the abdominal integument of *A. mellifera*, but not in the integument of the other two bee species. The genes *knk* and *Rtv* are both involved in cuticle stabilization in *Drosophila* [75]. In *T. castaneum*, *Rtv* activity is essential for localization of the Knk protein, facilitating its transport to the cuticle [76, 77]. The co-expression of *Rtv* and *knk* in *A. mellifera*, as shown in the reconstructed co-expression network, supports interaction of their respective products, as verified in *T. castaneum*. We also found *knk* transcripts in *C. analis* integument transcriptome, but not in *F. varia*. *Rtv* transcripts were not detected in the integument of these two bee species.

### Genes encoding desaturases and elongases potentially involved in CHC biosynthesis

CHC biosynthesis occurs in the epidermis-associated oenocytes [20] through biosynthetic pathways where desaturase and elongase enzymes have essential roles. Previously, we characterized the gene expression profiles of six desaturases and ten elongases in the developing integument of *A. mellifera* [23]. Our RNA-seq data confirmed these findings, besides identifying three more desaturase genes and other four genes encoding elongases potentially involved in CHC biosynthesis for deposition in the cuticular envelope. For *A. mellifera*, *F. varia* and *C. analis*, a higher proportion of the differentially expressed desaturase and elongase genes showed increased expression in the adults (Ne and/or Fg phases), and only for the eusocial species there were genes more expressed in the pharate-adults (Pbm phase). Among the desaturase and elongase genes, we highlight the expression profiles of *Desat-GB40659*, *Elo-GB54401*, *Elo-GB54302*, *Elo-GB45596* and *Elo-GB46038* orthologs, all showing positive correlation exclusively between the eusocial species.

### Genes of the ecdysone signaling cascade regulating cuticle formation and ecdysis in the integument

We detected in the integument the expression of genes that are part of the signaling cascade underlying insect molting and ecdysis, such as *EcR*, *Usp*, *E74*, *E75*, *FTZ-F1*, *CCAP*, *CCAPR* (*Crustacean Cardioactive Peptide Receptor*), *Eth* (*Ecdysis triggering hormone*), *Ethr*, and *Eh* [78]. Importantly, transcripts for these regulators were also detected in greater or lesser levels after ecdysis, in the integument of adult bees. *Usp*, which together with *EcR* forms the nuclear receptor complex that binds 20E and regulates the expression of a cascade of ecdysone-responsive genes, showed the higher expression in *A. mellifera* foragers. This is here tentatively related to the elevated JH titer at this phase of *A. mellifera* worker life [41] once *Usp* also has been proposed as a mediator of JH action [79].

*CCAP*, *hairy*, *mirr*, and *Kr-h1* in *A. mellifera*, *CCAP*, *Kr-h1*, *and Met* (*Methoprene-tolerant*) in *F. varia*, and *Kr-h1*, *E75*, *EcR, hairy*, and *mirr* in *C. analis* showed increased expression levels at the Ne and/or Fg phases. The roles of theses genes in adult bees, evidently dissociated from the molting events and metamorphosis, are yet to be determined. *Kr-h1* is a direct JH-response gene. *Met*, the JH receptor, has roles in the crosstalk of JH and 20E signaling pathways, which are critical in the regulation of insect metamorphosis [80].Since*Met*, and also *hairy*, mediate the action of JH on gene regulation [81], they certainly are needed in adult bees where JH has important physiological roles. The *mirr*geneencodesahomeodomaintranscriptionfactorwithrolesin*Drosophila*oogenesis[82]. Toour knowledge,itsroleintheintegumenthasnotyetbenstudied.

Some of the identified regulatory genes have been described as playing roles in cuticular melanization, as an example, the *Abdominal B* (*Abd-B*) Hox gene, which regulates *yellow* in the pigmentation/sclerotization pathway in *Drosophila* [83]. *Hairy*, which is a pair-rule gene in *Drosophila* embryos [84], may be involved in the polarity of abdominal segment melanization. The heterodimeric neuropeptide bursicon, composed by the gene products Bursα and Bursβ, is responsible for the regulation of the laccase2-encoding gene, and is crucial for the melanization/sclerotization of the newly formed cuticle [85, 86]. Bursicon interacts with the target tissue through its receptor, the product of the *rickets* gene, whose transcripts were also identified in our RNA-seq analysis of the integument of the three bee species.

### Searching for clues linking cuticle maturation heterochrony to eusocial or solitary life styles in the RNA-seq analysis

Our RNA-seq analyses were used to discover active genes in the integument of three bee species and, in addition, we looked for differences in gene expression profiles that could be linked to the heterochronic cuticle maturation dependent on the social/solitary ways of life. The following main findings highlighted differences in integument gene expression distinguishing the eusocial *A. mellifera* and *F. varia* from the solitary *C. analis*: **(a)** In contrast to the eusocial species, a smaller proportion of the genes differentially expressed in the integument was upregulated in *C. analis* foragers in comparison to the newly emerged bees, which is consistent with the cuticle of the solitary bee reaching maturity already at the emergence time; **(b)** The GO analysis including all the integument genes displaying orthology relationship with *Drosophila* genes highlighted functional categories that were mainly shared by both eusocial species in detriment of the solitary *C. analis*; **(c)** The Euclidean distance analysis based on the set of differentially expressed genes clearly separated the *A. mellifera* foragers from the newly-emerged, whereas in *C. analis* these bee groups were clustered, thus suggesting a greater similarity between the integument of newly-emerged and foragers of *C. analis*; **(d**) In contrast to the eusocial species, most of the genes for melanization/sclerotization, genes encoding RR1, RR2, or non-RR structural proteins, and also regulatory genes, did not show significant expression level variations in *C. analis*. Such differential fluctuation in transcript levels during development may have possibly contributed to the molecular heterochrony of cuticle maturation associated with bee life style. In addition, consistent with the comparatively earlier cuticle maturation process in the solitary bee, we found a higher proportion of CHC biosynthesis-related genes (desaturase and elongase genes) with significantly increased expression levels at the emergence (Ne phase) of *C. analis* in comparison to the eusocial bees. Furthermore, correlation analysis showed that a fraction of cuticle-related genes displayed congruent expression profiles between the eusocial species, but not with the solitary one; **(e)** By superposing the integument gene co-expression networks constructed for the three bee species, we found common interactions for the eusocial species, which were not seem when we compared these species with the solitary one.

Taken together, the comparative approach of the RNA-seq data highlighted suitable gene expression signatures related to adult cuticle formation and maturation in the bee species, in addition of revealing differences in gene expression that may possibly be involved in cuticle maturation heterochrony. Yet, this process may have entailed changes in the expression profiles of regulators of molting and metamorphosis.

### Abdominal adult cuticle deposition timing and its ultrastructure exhibit marked differences between the bee species

Cuticle ultrastructure and thickness did not significantly vary between the pharate adults (Pbm phase), newly-emerged (Ne) and foragers (Fg) of *F. varia*, as evidenced by TEM analysis. This was an unexpected result, considering that at the emergence time, *F. varia* workers visibly show an immature cuticle, i.e., incompletely pigmented and sclerotized. Therefore, the evident intensification of cuticular pigmentation and sclerotization in *F. varia* in the subsequent days after the emergence, which is necessary for flight and task performances outside the nest, do not imply in changes in abdominal cuticle thickness. It is possible, however, that thickness measurements taken from cuticular regions other than the abdominal, could evidence a different result, considering that regions of the insect body may diverge in the number of cuticle layers [87] and, consequently, in cuticle thickness.

In contrast, in *A. mellifera*, cuticle deposition is extended through the initial adult stage. Only in the honeybee we could identify post-ecdysially-deposited cuticle layers. Both, the pre-and post-ecdysially-deposited cuticle layers, or laminae, form the procuticle, which corresponds to the largest portion of the cuticle in insects in general. The term exocuticle has been used synonymously with pre-ecdysial cuticle, whereas those layers deposited post-ecdysially form the endocuticle. However, there is some divergence concerning these concepts [88]. In beetles, for example, up to three endocuticle layers are already present in specific areas of the body surface at the time of the adult ecdysis [87]. In *Sarcophaga bullata* flies, deposition of endocuticle occurs before the adult ecdysis [89].

As expected, the solitary bees, *C. analis* and *T. diversipes*, and even the primitively eusocial *B. brasiliensis* and the facultatively eusocial *E. cordata*, showed a fully deposited cuticle as soon as they emerge, and newly-emerged and forager bees in each of these species displayed similar cuticle ultrastructure, pigmentation and sclerotization. The rapid cuticle maturation in *E. cordata* is consistent with its nesting biology and social structure. *E. cordata* nests are founded by a single female that build up until ten brood cells. The offspring will leave the nest immediately after the emergence for founding new nests. However, daughters may return to the maternal nest, thus resulting in a facultatively social organization with a dominant female (the mother) and its subordinate daughters. There are also nests formed by sisters’ females or even by unrelated females, the oldest one showing dominance over the youngest. The dominant female produces all the offspring and rarely leaves the nest, whereas the subordinates assume the tasks of nest provisioning and maintenance, and they also lay trophic eggs that are eaten by the dominant [90–92]. Such female associations may have preceded the highly eusocial way of life [93]. Therefore, in *E. cordata*, as well as in the truly solitary bees, *C. analis* and *T. diversipes*, rapid cuticle maturation is the condition for the immediate exit from the nest after emergence.

This situation is somewhat diverse for the primitively eusocial *Bombus*. In *B. brasiliensis*, as demonstrated here, the final adult cuticle ultrastructure and thickness are achieved at the emergence. This would allow the workers start foraging soon, as reported for *B. atratus* workers that may leave the nest as soon as at the emergence day (0 day). However, workers of this species may start foraging later, at the age of 10-20 days [94], thus similar to the eusocial bees. Moreover, younger workers in the genus *Bombus* have, in general, incompletely pigmented cuticle and hairs, denoting that cuticle maturity was not yet completely achieved. Such characteristics that seem intermediary to the eusocial and solitary condition may be inherent to the primitively eusocial species, but this requires further investigation. Studies correlating the grade of cuticle pigmentation with the age of starting foraging among primitively eusocial bee species should clarify this issue.

Our TEM analysis and thickness measurements showed that in the same abdominal segment, cuticle ultrastructure greatly differs between the bee species, not only in the number of the adjacently arranged chitin/protein sheets (laminae), but also in the morphology of the most superficial layers. Except for *F. varia*, these results are consistent with a cuticle development timing adapted to the life style, as observed for the highly eusocial *A. mellifera*, the facultatively eusocial *E. cordata*, the primitively eusocial *B. brasiliensis*, and the solitary *C. analis* and *T. diversipes* bees.

Considering that the timing of cuticle deposition is peculiar to bee species, and that cuticle deposition rhythm in *Drosophila* is regulated by a peripheral circadian oscillator in the epidermal cells, which requires the expression of the clock genes *Per*, *Tim2*, *Cyc*, and *Clk* [39], and also that a *Cry* clock gene regulates the rhythm of cuticle deposition in the bean bug *Riptortus pedestris* [40], we compared the expression of seven circadian rhythm genes (*Per*, *Tim2*, *Cyc, Clk*, *Cry, Vri and Pdp1*) in the developing integument of *A. mellifera*, *F. varia* and *C. analis*. Consistent with the differences in the timing of cuticle deposition, the expression profiles of *Clk* in *A. mellifera*, *F. varia*, and *C. analis* were negatively or non-correlated. Similarly, the expression profiles of *Cry* in *A. mellifera* and *C. analis* were negatively correlated (*Cry* was not identified in *F. varia*), as well as the expression of *Per*. It is likely that *AmPer* has roles in adult cuticle organization. Interaction of *AmPer* and other genes involved in cuticle formation was specifically observed in *A. mellifera*, whose sequenced genome gives more support for gene co-expression network reconstruction. In *A. mellifera*, *Per* was co-expressed with the *knk* gene, which in *T. castaneum* was associated with stabilization of the cuticular laminae [95]. Both genes were co-expressed with structural cuticular protein genes such as *AmCpap3-a*, *AmTwdl*(*Grp*-Glycine-rich protein), *AmUnCPR-RR2-2*, *AmCPR26*, *Am49Ah-like* and *AmSgAbd2-like*, and also with *Amyellow-y*, a gene in the yellow family, involved in cuticle pigmentation [96]. The expression profiles of another clock gene, *Tim2*, were positively correlated between the eusocial species, with a marked decrease in expression levels at the emergence, suggesting roles in the final step of adult cuticle formation in these bees. The *Pdp1* gene encodes a basic leucine zipper transcription factor and is expressed at high levels in the epidermis and other tissues of *Drosophila* embryos. *Pdp1* is an essential clock gene linked to the circadian rhythm. It is a regulator of *Clk* and other clock genes, such as *Tim*, *Per*, and *Pdf* (Pigment dispersing factor), a neuropeptide controlling circadian behavioral rhythms [97, 98]. *Pdp1* seems an important gene in the *C. analis* integument since it is connected with nine structural cuticle protein genes, three chitin-related genes, and two desaturase encoding genes in the co-expression network. However, it is significantly more expressed after the emergence, when the cuticle of the solitary bee is already formed, thus virtually excluding a role in cuticle laminae deposition rhythm. Differently from *C. analis*, *Pdp1* was not co-expressed in the networks reconstructed with the *A. mellifera* and *F. varia* genes involved in cuticle formation and maturation. Some of the cuticular genes were also co-expressed with *Cyc* in the integument of *A. mellifera* and *C. analis*.

### Cuticular n-alkanes as markers of cuticle maturity in bees

N-alkanes are structural lipids in the insect cuticle [99, 17], where they compose the envelope [100]. The absolute quantities of n-alkanes and unsaturated CHCs were significantly higher in the foragers than in the earlier developmental phases of the eusocial *A. mellifera* and *F. varia* species. The n-alkanes detected in higher proportions in *A. mellifera* foragers than in the newly-emerged were C_23_, C_24_, C_25_, C_26_, C_27_, C_29_, C_31_, and C_33_, the C_25_ and C_27_ n-alkanes presenting the highest proportions. The analysis of the individual CHC peaks obtained from *F. varia* also showed higher proportions of C_27_ and C_29_, besides a higher proportion of C_22_, in the foragers. All these n-alkanes, except C_27_ and C_33_, were also proportionally increased in foragers than in newly-emerged bees of the eusocial *Melipona marginata* [101]. These data are consistent with previous reports on higher levels of n-alkanes in *A. mellifera* foragers [102] and in foragers of an ant species, *Pogonomyrmex barbatus* [103]. In contrast, the proportions and absolute quantities of n-alkanes did not differentiate foragers from the newly-emerged in *C. analis*. Together, these findings may be interpreted as the solitary bee displaying an accelerated process of cuticle maturation in comparison to the eusocial ones. N-alkanes may be markers of cuticle structure maturation. Long-chain alkanes are thought to increase cuticle waterproofing [104, 103], suggesting that this essential ability for the performance of extra-nidal activities was acquired earlier in the development of *C. analis*. At the adult emergence, the solitary bee already has the chemical profile needed for a prompt interaction with the environment outside the nest. Consistently, the levels of n-alkanes also did not significantly differ between young and old females of the solitary leafcutter bee species, *Megachile rotundata* [105].

## Conclusions

Using RNA-seq analysis of the integument of two eusocial bee species, *A. mellifera* and *F. varia*, and a solitary bee, *C. analis*, we identified genes involved in cuticle (exoskeleton) formation and maturation. The expression profiles of these genes were determined at three developmental time points corresponding to adult cuticle deposition/differentiation at the pharate-adult stage, newly-ecdysed cuticle, and fully developed cuticle of forager bees. TEM analysis of the cuticle at these time points, including other bee species, and CHC profiles determination were performed in addition to the transcriptome analysis. Together, these experimental approaches provided novel data on integument developmet. We also searchedforcluesinintegumentgeneexpresion,structure,andCHCprofilesthat could be consistent with the premise that eusociality might have entailed heterochronic changes in cuticle development, resulting in faster cuticle maturation in the solitary bee, thus allowing flight and forager activities immediately after emergence, and in slow cuticle maturation in the eusocial bees, which benefit from the protected nest environment for a period of time after the emergence. This study expands our understanding on the molecular biology and structure of the developing integument, besides highlighting differences in the process of cuticle maturation related to the eusocial/solitary behaviors.

## Materials and Methods

### Sample collection

We collected workers of *A. mellifera* (Africanized) and *F. varia* from colonies maintained in the Experimental Apiary of the Faculdade de Medicina de Ribeirão Preto, Universidade de São Paulo, Ribeirão Preto, SP, Brazil. Trap-nests to collect samples of the solitary species *C. analis* and *T. diversipes* were made [106] and placed in the Experimental Apiary area. Additional bee species (*B. brasiliensis*, *E. cordata* and *T. diversipes*) were obtained from donations (see acknowledgments section).

We used females from three developmental phases: pharate adults in process of cuticle pigmentation (Pbm phase), newly emerged (Ne) adults and foragers. Carrying pollen bees from the solitary and social species, and building-nest females from the solitary species, were identified as foragers (Fg). The *B. brasiliensis*, *E. cordata* and *T. diversipes* species were exclusively used for cuticle morphology studies through TEM. In this case, we used the Ne and Fg phases.

### RNA extraction and sequencing

For each developmental phase (Pbm, Ne and Fg) of *A. mellifera* and *F. varia*, we prepared three independent samples, each made with five abdominal integuments. For the corresponding developmental phases of *C. analis*, we prepared three independent samples, each containing three abdominal integuments. The RNA extractions were made using TRIzol^®^ reagent (Invitrogen) following manufacturer's instruction. The extracted RNAs (2 µg/per sample) were sent to a facility (Laboratório Central de Tecnologias de Alto Desempenho em Ciências da Vida, Universidade Estadual de Campinas, Campinas, Brazil) to access sample quality through a 2100 Bioanalyzer and for library preparation (TruSeq™ RNA - Illumina^®^) and RNA sequencing in an Illumina HiSeq 2500 equipment (paired-end reads, 2 × 100 bp read length). We obtained an average of 30 million reads per sample, with 90% of the bases showing quality scores > Q30. The RNA-seq data is deposited at the National Center for Biotechnology Information (NCBI) database under the BioProject ID PRJNA490324.

### Adapters trimming and quality check

We used the software Scythe v. 0.991 (https://github.com/vsbuffalo/scythe) for trimming 3' standard Illumina adapter sequence. We followed a Cutadapt v. 1.4.1 [107] trimming at 5’ ends of the reads, and we filtered reads with Phred quality > 20. The trimmed sequences were filtered using the software PRINSEQ-lite v. 0.20.3 [108] and sequence quality was evaluated through the software FastQC v. 0.11.2 [109].

### Transcriptome assembly and gene expression

We aligned the high quality reads from *A. mellifera* against its genome v. 4.5 using the software TopHat v. 2.0.9 [110]. The *A. mellifera* aligned sequences were quantified and the developmental phases compared using the software Cufflinks v. 2.1.1 [111]. The extension Cuffmerge integrated the reads to the mapping results and the tool Cuffdiff checked the expression levels for each sample and the significance of comparisons. CuffmeRbund R package v. 2.8.2 allowed us to access all this information [112].

For *F. varia* and *C. analis*, we used the software Trinity (trinityrnaseq_r2014717) [113, 114]. The N50 contig length (smallest contig length for which the sum of fewest contigs corresponds to 50% or more of the assembly [115]) of all transcripts of *F. varia* was 2372 and of *C. analis* was 2440. Orthology search was performed through the software InParanoid 8 [116]. We only accepted those transcripts with higher similarity with *A. mellifera* than to *Drosophila melanogaster*. Statistical evaluation of these data was done with the R software v. 3.1.2, using the packages R DESeq2 v. 1.6.3 [117] and edgeR v. 3.8.6 [118]. We considered as differentially expressed between developmental phases, those contigs with significant results for both R packages.

All heat maps were designed using the function heatmap.2 from gplots R package [119]. For all groups of genes, we measured the clustering potential of the samples for each phase and species. For this approach, we used the R package pvclust v. 1.3.2 [120] based on correlation distances, with a complete linkage method, and 10,000 bootstrap replication. We used unbiased p values (AU) and bootstrap values as measurements of clusters’ significance. Clusters showing AU > 95% were considered statistically significant [120].

### Molecular and functional characterization of differentially expressed genes

For the analysis of gene expression in *A. mellifera*, we filtered the differentially expressed genes using the following thresholds: q-value < 0.05; Log 2 Fold Change ≤ −1 or ≥ 1 and Fragments Per Kilobase of transcript per Million mapped reads (FPKM) ≥ 5. In the case of the other two bee species, *F. varia* and *C. analis*, we used the parameters cited in the previous section. For the Gene Ontology (GO) enrichment analysis we used the *A. mellifera* gene IDs to look for *D. melanogaster* ortologues in Fly Base, through the support of the online softwares g:Profiler (http://biit.cs.ut.ee/gprofiler/gorth.cgi), and g:Orth function [121, 122]. The same was done for *F. varia* and *C. analis* but using *A. mellifera* ortologues. We filtered the *Drosophila* IDs to avoid ID repetition and used them to generate an input list for the software DAVID v. 6.7 (http://david.abcc.ncifcrf.gov) [123, 124], used to perform the Gene Ontology analysis. The annotated functions belonged to Biological Process (BioP), Cellular Components (CC) and Molecular Function (MF) categories. Structural cuticular protein encoding genes were classified in accordance with the software CutProtFam-Pred (http://aias.biol.uoa.gr/CutProtFam-Pred/home.php) [125]. Venn diagrams were plotted with the online version of the software jvenn [126] (<http://bioinfo.genotoul.fr/jvenn/example.html>).

### Transcription factor binding sites search with TRANSFAC^®^

Transcription factors whose binding sites could be enriched in specific groups of *A. mellifera* gene models were searched using TRANSFAC^®^ [127] against insects database. For this enrichment analysis, we searched the 5’ UTR regions covering −3,000 bases relative to the transcription start sites of the genes involved in sclerotization/melanization processes, chitin metabolism, CHC biosynthetic pathways, regulation of cuticle formation and maturation, circadian rhythm, and non-melanization pigmentation pathways (see gene IDs S1 File). We excluded the genes with 5’ UTR < 500 bases. We used the *A. mellifera* genes in this analysis once it is the only among the species here studied with an available reference genome. For the transcription factor FTZ-F1 binding sites, we used the TRANSFAC database (DROME$FTZF1_01, DROME$FTZF1_02, and DROME$FTZF1_03 from *D. melanogaster*, and BOMMO$CPR92_01, and BOMMO$CPR92_02 from *B. mori*) to generate a positional matrix. Here, we only highlighted those transcription factor binding sites (TFBS), which could be relevant for insect cuticle formation and maturation (see **Discussion section**).

### Gene co-expression networks

The networks were plotted based on the correlation of gene expression in the integument of the analyzed bee species. We used the software Cytoscape v. 3.3.0 [128] – for Linux, and its plugin ExpressionCorrelation App v. 1.1.0. We only accepted correlations ≥ +0.95 or ≤ −0.95 and p ≤ 0.05.

### Transmission electron microscopy (TEM)

We dissected the integument from the right anterior region of the third abdominal tergite of the studied bee species. The ultrastructure of the integument was compared between species and between the developmental phases (Pbm, Ne and Fg) using 11 *A. mellifera* integument pieces (4 from Pbm, 3 from Ne, and 4 from Fg phases), 9 integuments from *F. varia* (3 Pbm, 3 Ne, 3 Fg), 11 integuments from *C. analis* (4 Pbm, 4 Ne, 3 Fg), 4 integuments from *B. brasiliensis* (2 Ne, 2 Fg), 6 integuments from *E. cordata* (3 Ne, 3 Fg), and 6 integuments from *T. diversipes* (3 Ne, and 3 Fg). The Ne and Fg phases of *brasiliensis* were recognized based on the grade of body pigmentation criterion (less intense in the Ne bees) and for the Fg phase we also examined the wings in the search for erosion signals that could indicate intense foraging activity. The integument samples were fixed in 5% glutaraldehyde in cacodylate buffer 0.1 M, pH 7.2, during 2 h under shaking, washed 3X in the cacodylate buffer, fixed in osmium tetroxide 1% diluted in phosphate buffer 0.1M, pH 7.2, dehydrated in acetone and propylene oxide and embedded in resin. We used uranyl acetate for enhancing image contrast. The ultrathin sections were examined in a Jeol-Jem-100cx-II Electron Microscope and the software ImageJ v. 10.2 [129] was used to measure integument thickness. Measurements were compared among the developmental phases and species using Analysis of Variance (ANOVA) associated with the Tukey's Honestly Significance Difference (Tukey's HSD) post hoc test in R software v. 3.1.2, except for the *A. mellifera* and *B. brasiliensis* data. As for *A. mellifera* the data did not present a normal distribution (Shapiro-Wilk normality test) we used the Kruskal-Wallis test associated with the *post hoc* Conover-Iman test and Bonferroni correction, and for *B. brasiliensis* we used the Student’s t-test [130].

### Cuticular hydrocarbon profiles

The quantification of CHCs was based on their peak area in each chromatogram. For the analysis of relative peak area, we collected 15 bees per developmental phase for each of the highly eusocial species, while for *C. analis* we obtained four Pbm-staged bees, seven Ne and seven Fg bees. The *F. varia* hind legs were removed to avoid resin contamination. Except for this species, we bathed each sample in 1.5 ml of n-hexane 95% (Mallinckrodt Chemicals) for 1 min and 30 s to extract the CHCs. Due to the small size of *F. varia* species, we used 500 μl of n-hexane to extract the CHCs [131]. The extracts were dried under N2 flow and resuspended in 160 μl of n-hexane (100 μl for *F. varia* extracts) before running the analysis. CHC identification was made in a Gas Chromatograph / Mass Spectrometer (GC-MS) system (Shimadzu GCMS model QP2010), equipped with a 30 m DB-5MS column using helium as the carrier gas (1 ml/min), through electronic ionization (EI) mode. CHC relative quantification and normalization of the peak areas were performed following Falcón *et al.* [23] description. We also compared each developmental phase considering the classes of CHC (n-alkanes, unsaturated, and branched alkanes), repeating the normalization process for each case.

In order to verify differences between the developmental phases and between the bee species, we performed a clustering approach as described in the section **Transcriptome assembly and gene expression**, but using the Euclidean distance instead of the correlation distance. We also verified the compounds that better explained the detected differences. With this purpose, we performed a Principal Component Analysis (PCA) using the R software. The variation of each CHC peak area between the developmental phases was accessed through a Tukey's HSD test in R software.

Additionally, we calculated the absolute quantities of n-alkanes and unsaturated CHC per bee for the species *A. mellifera*, *F. varia*, and *C. analis*. An analytical curve [132] was built to establish the correlation between the quantities of the used standards and the CHCs. It is described by the equation: y=ax +b, where y is the known amount of the standard, × is the quantity of the unknown CHC, a is the peak area of the standard, and b is the area of the intercepted background. We prepared the curve based on the n-alkanes standards C_23_, C_25_, and C_32_ (Alltech Corporation) and using 1.25, 2.5, 5, 10, 15 and 20 μg/ml of each alkane. To the standard solutions and to each sample, we added 100 μl of the internal standard α-colestane (6.25 μg/μl) (Sigma-Aldrich) under the previously cited chromatography conditions. The values of the correlation curve (R) between CHCs and standards were ≥ 0.99. After curve preparation, CHCs were quantified in three independent samples, each prepared with individual bees, for each developmental phase and species. Due to the reduced body size of *F. varia*, we used pools of three bees for each one of the three independent samples per developmental phase and the obtained CHC values were corrected accordingly. To calculate the concentrations of compounds up to C_24_, we used the curve of C_23_; for compounds from C_25_ up to C_29_, we used the C_25_ curve; and for compounds larger than C_29_ up to C_35_ we used the curve of C_32_. We followed an ANOVA associated with the *post hoc* Tukey’s HSD test to compare the absolute quantity of each CHC (n-alkanes and unsaturated) between developmental phases. The absolute quantities of the *F. varia* unsaturated compounds were not calculated once their quantities are very low.

## Acknowledgments

We thank to Sidnei Matheus, Solange Augusto, Gabriela Freiria and Anete Lourenço by the samples provided. This work was supported by São Paulo Research Foundation (FAPESP): Grants to MMGB (2014/13136-0), ZLPS (2011/03171-5), and NPL (2014/50265-3). Fellowships to: TF (2012/24284-5) – during PhD, MEN (2012/09108-6). And later, fellowship from Programa Nacional de Pós-Doutorado (PNPD) CAPES/HCPA to TF (Process number: 88887.160608/2017-00). This study was financed in part by the Coordenação de Aperfeiçoamento de Pessoal de Nível Superior - Brasil (CAPES) - Finance Code 001.

## Supporting information

**S1 Fig.** Correlation heatmaps based on the RNA-seq data obtained from the integument of the pharate adults (Pbm), newly-emerged (Ne) and foragers (Fg) of the three bee species: (A) *A. mellifera*, (B) *F. varia* and (C) *C. analis*. The numbers 1, 2 and 3 following the Pbm, Ne, and Fg abbreviations indicate the independent samples of each developmental phase.

**S2 Fig. Gene co-expression networks in the integument of *A. mellifera* for adult cuticle formation and maturation.** The genes are indicated in the nodes, and the edges represent significant correlation among genes.

**S3 Fig. Gene co-expression networks in the integument of *F. varia* for adult cuticle formation and maturation.** The genes are indicated in the nodes, and the edges represent significant correlation among genes.

**S4 Fig. Gene co-expression networks in the integument of *C. analis* for adult cuticle formation and maturation.** The genes are indicated in the nodes, and the edges represent significant correlation among genes.

**S5 Fig. Distances between the developmental phases of *A. mellifera*, *F. varia* and *C. analis* based on Euclidean distance analysis of total CHCs, n-alkanes, unsaturated CHCs, and branched CHCs relative quantifications.** Red boxes indicate significant clusters with 95% of confidence. Arrows indicate significantly supported clusters. AU clusters’ support (red values); BP bootstrap support (green values); Branches’ edges (gray values). Pharate-adults (Pbm), newly emerged (Ne), and forager (Fg) bees.

**S1 Table. Ortholog genes displaying significantly correlated expression profiles.** Comparisons of gene expression levels through the pharate-adult (Pbm), newly-emerged (Ne) and forager (Fg) developmental phases of *A. mellifera*, *F. varia*, and *C. analis*. Blue: significant positive correlation. Red: significant negative correlation. (-): undetectable gene expression.

**S2 Table – Number of genes encoding the different classes of structural cuticular proteins in hymenopterans.**

**S1 File. Genes identified in the RNA-seq analysis of the integument of *A. mellifera*, *F. varia* and *C. analis*.**

**S2 File. Genes upregulated in the comparisons of the developmental phases of each bee species, *A. mellifera*, *F. varia* and *C. analis*.** Pharate-adult (Pbm), newly-emerged (Ne) and forager (Fg) developmental phases. The bee species and developmental phases compared are specified in the inferior margin of each table in this File. For example: Amel_Pbm>Ne means the list of *A. mellifera* genes upregulated in Pbm in comparison to Ne.

**S3 File. Gene Ontology (GO) functional analysis of the differentially expressed genes.** The bee species and developmental phases compared are specified in the inferior margin of each table in this File. For example: Amel_Pbm_Ne>Fg_ID means the bees’ gene IDs and Fly Base gene IDs from higher expressed genes at younger developmental phases; and Amel_Ne_Fg>Pbm_GO means the Gene Ontology of the higher expressed genes at older developmental phase based on their fly orthologues IDs.

**S4 File. Cuticular hydrocarbon (CHC) profiles determined for the developmental phases of *A. mellifera*, *F. varia*, and *C. analis*, variable contribution (total and per comparison) and mass quantification.** Pharate-adult (Pbm), newly-emerged (Ne) and forager (Fg) developmental phases.

